# Green nanoparticles of Ag, SiO_2_, and CeO_2_: strategy for controlling the PVY–ToBRFV coinfection in bell pepper

**DOI:** 10.64898/2025.12.30.696975

**Authors:** Ubilfrido Vasquez-Gutierrez, Sonia N. Ramírez-Barrón, Gustavo A. Frias-Treviño, Luis A. Aguirre-Uribe, Agustín Hernández-Juárez

## Abstract

Bell pepper (*Capsicum annuum* L.), a crop of global importance, is threatened by mixed viral infections, such as Potyvirus yituburosum (PVY) and Tobamovirus fructirugosum (ToBRFV). This study evaluated the antiviral potential of green-synthesized silver (Ag), silicon dioxide (SiO_2_), and cerium dioxide (CeO_2_) nanoparticles (NPs) against these viruses in seeds and seedlings. NPs were synthesized using aqueous walnut shell extracts. The nanoparticles were characterized by EDX and TEM. In two experimental phases, bell pepper seeds were treated with different concentrations of nanoparticles (50–1000 mg L^−1^) and subsequently exposed to PVY, ToBRFV, and both. Viral transmission was assessed by qELISA in seeds and seedlings. Seedling development, germination rates, and virus incidence were measured; phytotoxicity (CF_50_), antiviral efficacy (IC_50_/IC_90_), and the selectivity index (SI) were calculated. The results showed a significant reduction in viral load and inhibition of PVY and ToBRFV transmission in both single and mixed infections, especially at doses ≥400 mg L^−1^. CeO_2_ nanoparticles exhibited the highest selectivity indices (SI > 3000) and no phytotoxicity. Ag and SiO_2_ nanoparticles also exhibited antiviral activity, but with slightly higher CF_50_ values. In the greenhouse, seedlings grown from seeds treated with nanoparticles showed a reduction in viral symptoms and an increase in SPAD index and height. In conclusion, bio-nanoparticles, especially CeO_2_, are promising antiviral agents against PVY and ToBRFV in pepper cultivation. Their high sensitivity and low toxicity make them good candidates for the sustainable management of plant viruses, justifying further field-scale evaluation.

## Introduction

Bell peppers (*Capsicum annuum*), known as sweet peppers for their rich nutritional content, are a crop of global importance in 2024 (Devi et al., 2021). They are low in calories and rich in essential vitamins and antioxidants, including vitamins A and C, potassium, folate, and fiber (Ahmed et al., 2024). In 2022, global production of fresh bell peppers was approximately 52.14 billion kilograms (kg), with China as the leading producer at 16.81 billion kg. 81 billion kg, with Mexico and Indonesia among the top producers, at 3.11 and 3 billion kilograms, respectively (Tridge, 2025). In Mexico, the producing states include Sinaloa, Chihuahua, Sonora, Zacatecas, and Jalisco; Sinaloa stands out as the main open-field pepper producer, with an annual output of around 166,000 tons, while Chihuahua has been recognized for its 23.6% share during the 2016–2020 period (SIAP, 2025). The growth in production underscores the importance of peppers in agriculture and global trade (Biswas et al., 2018); however, climate-induced yield losses and the emergence of new pests highlight the need for resilient agricultural practices (Zakir et al., 2024). Despite these challenges, a compound annual growth rate of 8% is expected between 2024 and 2031, reflecting their enduring importance in global agriculture and gastronomy (Cognitive Market Research, 2025).

Bell pepper plants are susceptible to a wide range of viral species, including cucumber mosaic virus (CMV), pepper mild mottle virus (PMMoV), potato virus Y (PVY), tobacco mosaic virus (TMV), pepper mottle virus (PepMoV), alfalfa mosaic virus (AMV), tomato spotted wilt virus (TSWV), and pepper leaf curl virus (PepLCV); most of these viruses are transmitted by insect vectors, seeds, or plant debris (Arogundade et al., 2020; Parisii et al., 2020). Infected plants exhibit symptoms such as mosaic or mottling patterns, chlorosis, growth retardation, leaf curling or distortion, and malformed or discolored fruits, all of which contribute to a significant reduction in plant vigor and yield (Ormeño et al., 2006; Makhsudova et al., 2024; Rodríguez-Román et al., 2023). Sometimes these plant syndromes result from the synergistic infection of two or more viral species, commonly referred to as mixed infections (Sánchez-Tovar et al., 2025). PepLCV can cause growth retardation, flower bud abortion, reduced pollen production, and losses of up to 90–100% (Dwivedi et al., 2024), especially in plants infected at early stages. Meanwhile, CMV reduces yield by more than 80% during severe outbreaks (Li et al., 2020). Research on mixed infections with PVY and ToBRFV remains limited, but parallels have been drawn with well-documented cases of mixed viruses in solanaceous crops, underscoring their serious implications for plant health and disease management. In tomato, for example, co-infection with ToBRFV and PepMV has been shown to intensify symptoms (immature fruits with scarring and pronounced foliar symptoms), suggesting possible synergistic interactions that increase disease severity (Yilmaz et al., 2023). Observations also indicate that ToBRFV can increase the viral titer of PepMV (Klap et al., 2020; Yilmaz et al., 2023), highlighting the risk of mixed infections. Although no direct reports of PVY and ToBRFV coinfection in pepper or tomato have been documented to date, the high adaptability and persistence of both viruses raise valid concerns about their potential interaction and cumulative impact on crops (Fidan et al., 2024; Yilmaz et al., 2023).

Given these threats to bell pepper crops, it is essential to investigate management strategies such as strict hygiene protocols, heat treatment of seeds, and reliable diagnostics to prevent viral spread (Vasquez-Gutierrez et al., 2025), especially since ToBRFV can survive on equipment for extended periods (Vasquez-Gutierrez et al., 2024a) and PVY persists in fields via aphid vectors; even potato tuber seeds harvested from PVY-infected plants grown in open fields can cause secondary PVY infections and result in production losses of up to 30% (De Santiago Meza et al., 2025). The nonspecific and overlapping nature of the symptoms, combined with the variety of transmission routes, makes early detection and differentiation of these viral infections extremely difficult, often resulting in delayed or inadequate treatment responses (Tatineni & Hein et al., 2023). Nanomaterials, particularly metallic nanoparticles such as silver (Ag), copper/copper oxide (Cu/CuO), and zinc oxide (ZnO), are emerging as promising tools for the diagnosis, prevention, and mitigation of plant viral diseases because they can act directly on viral particles, interfere with viral entry and replication, and/or stimulate the host’s antiviral defenses (Al-Askar et al., 2023; Liu et al., 2024).

Silver nanoparticles (AgNPs) have been repeatedly reported to reduce viral load and delay replication in infected plants, and several studies show antiviral effects against tobamoviruses and other plant viruses (Shahzadi et al., 2025); proposed mechanisms include direct binding to virions, disruption of viral proteins/nucleic acids, and generation of reactive oxygen species that reduce infectivity (Liu et al., 2024; Warghane et al., 2024). The development and implementation of these methods are essential to protect pepper and tomato production against potentially devastating mixed-virus outbreaks, highlighting the devastating economic impact of viral diseases on pepper production (Carrillo-Lopez et al., 2024).

Cu/CuO NPs exhibit antiviral activity and practical broad-spectrum antimicrobial advantages; recent studies of Cu NPs fragment TMV particles in vitro and reduce infectivity in plants (Liu et al., 2024), supporting their potential to protect solanaceous crops such as pepper. The “green” synthesis of metal nanoparticles produces stable, biofunctional nanomaterials using plant extracts or agricultural waste as reducing/stabilizing agents, which often reduces toxic reagents, lowers costs, and can add beneficial surface biomolecules that enhance biocompatibility and antiviral performance in foliar or seed treatments (Ahmad et al., 2022; Shahzadi et al., 2025). Despite the promising laboratory and greenhouse results, scaling up to field management is still limited: there are knowledge gaps regarding optimal dosages, application timing, effects on beneficial organisms, and long-term phytotoxicity, so integrated evaluation and well-designed field trials are essential before large-scale adoption (Carrillo-Lopez et al., 2024; Warghane et al., 2024). In this study, for the first time to our knowledge, the co-infection of ToBRFV and PVY in pepper plants is investigated, along with the effect of antiviral nanoparticles on mixed and individual infections from seeds to seedlings, and the impact on phytochemical compounds and radicals.

## 1. Materials and methods

### 1.1. Green synthesis and characterization of silver, silicon dioxide, and cerium oxide nanoparticles

#### Silver nanoparticles (AgNPs)

were synthesized in the Department of Basic Sciences at the Universidad Autónoma Agraria Antonio Narro in Saltillo, Coahuila, using the green synthesis technique of Neira-Vielma et al. (2022), employing an aqueous extract of pecan nut shells from Carya illinoinensis. The shells were ground in a ball mill (Fritsch, Pulverisette, Germany) to obtain a fine powder. One gram of shell powder was suspended in 200 mL of distilled water (pH 6.5) in a three-necked round-bottom flask connected to a reflux condenser and placed on a hot plate for 4 hours at 80 °C under magnetic stirring. For the synthesis of AgNPs, a solution of 1.3 g of silver nitrate (AgNO_3_) in 780 mL of distilled water (H_2_O) and 20 mL of walnut shell extract was prepared. The solution was placed in a three-neck round-bottom flask and stirred on a hot plate for two hours at 95 °C, then transferred to amber glass vials for refrigerated storage.

#### Silicon dioxide nanoparticles (SiO_2_)

a solution of 0.6 g of cetrimonium bromide (CTAB) was prepared. in 100 mL of H_2_O in a volumetric flask. 100 mL of ethanol, 200 mL of H_2_O, 50 mL of aqueous walnut shell extract, 4 mL of ammonium hydroxide, and 17.6 mL of tetraethyl orthosilicate were added to a three-necked round-bottom flask and placed on a stirring hot plate at 90 °C for 1 h. The resulting NPs solution was centrifuged at 3500 rpm for 3 min. The nanoparticle powder was placed in a drying oven (ICB-Oven) at 70 °C for 48 hours. At the end of the incubation period, the nanoparticle powders were removed and placed in a porcelain crucible inside a Type 1500 muffle furnace (Thermolyne, Thermo Fisher Scientific) at 500 °C for 1 hour. The nanoparticle powder was dissolved in a 50% anhydrous ethanol solution with distilled H_2_O under stirring, and then centrifuged at 3500 rpm. The organic phase was recovered and placed in tubes inside an oven for drying. Finally, the SiO_2_ NPs were homogenized and stored at room temperature.

Cerium oxide nanoparticles (CeO_2_): 1 g of cerium carbonate hydrate III (Sigma) was dissolved in 200 mL of ethanol and stabilized at pH 7.0. 200 mL of H_2_O, 50 mL of the aqueous walnut shell extract, and 4 mL of ammonium hydroxide were added to a round-bottom flask on a hot plate and stirred at 90 °C for 4 h. The resulting NPs solution was centrifuged at 3500 rpm for 3 min. The organic phase (NPs powders) was recovered and placed in a drying oven at 70 °C for 24 hours. The organic phase was recovered in a porcelain crucible and placed in a Type 1500 muffle furnace (Thermolyne, Thermo Fisher Scientific) at 500 °C for one hour. The NP granules were homogenized and stored at room temperature.

The characterization of the AgNPs was carried out by energy-dispersive X-ray analysis (EDX) for Ag and X-ray diffraction (XRD) for SiO_2_ and CeO_2_. To determine the crystalline domain size of the SiO_2_ nanoparticles, the Scherrer formula was used:

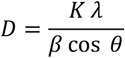

Where:

*D*= average crystallite size (nm).

*K*= shape constant (typically 0.9).

*λ*= Wavelength of X-ray radiation (for Cu Kα: 0.15406 nm).

*β*= Full width at half maximum (FWHM) of the peak corrected for instrumental broadening, in radians.

*θ*=angle 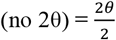, in radians.

To measure the full width at half maximum (FWHM) in the diffractogram. The peak was located (the maximum at 2θ = 22.4°).

The half-maximum was calculated, and the two points on the peak’s edge where the intensity was half-maximum were located; the left and right 2θ values were recorded. FWHM (in °) = 2*θ*_right_ − 2*θ*_left_. The points are located at 21.9° and 22.9°, FWHM = 1.0°. The instrumental correction was performed with the instrumental width (*β*_inst_),

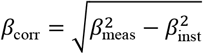

The conversion to radians was performed.

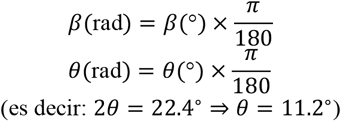

For cerium oxide (CeO_2_) nanoparticles, an estimation in degrees and Scherrer size was performed. CeO_2_ crystallizes in the fluorite-type cubic structure (space group Fm-3m), and its theoretical diffractogram is well tabulated. The characteristic angles for radiation Cu Kα (λ = 1.5406 Å). Subsequently, the pixels were converted to degrees. Automatic marker detection yielded unreliable results, so a fallback linear mapping of 0 → 80° across the plot’s internal width (0–80°) was used to convert pixels to degrees.

The following were used to estimate the crystallite size:

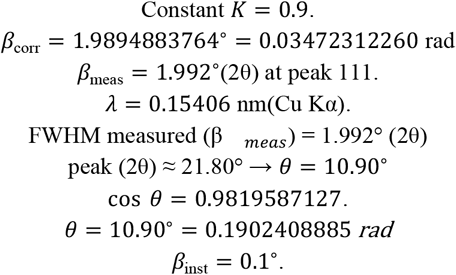

### 1.2. Isolation and preservation of ToBRFV and PVY in pepper plants

Plant samples exhibiting characteristic viral symptoms were collected from bell pepper farms in General Cepeda, Coahuila, Mexico (25°22′42.8″N 101°25′25.7″W); the leaves were individually analyzed and characterized serologically and molecularly. Viral purification of PVY and ToBRFV was performed by inoculating *Nicotiana longiflora* plants, employing the technique of Vásquez-Gutiérrez et al. (2024). For this purpose, a ToBRFV inoculum was prepared from symptomatic tissue of the collected samples, macerated (1:10, w/v) in phosphate-buffered saline (PBS) at pH 8 (0.01 M), and supplemented with celite (Sigma, Darmstadt, HE, Germany) as an abrasive. Using a swab soaked with 100 µL of infectious sap, the sap was spread onto the two apical leaflets of each plant. Subsequently, the plants were washed with PBS. Five replicates were used for each virus inoculated into plants. The observation of local and systemic symptoms was quantified 20 days post-inoculation (dpi). The same inoculation procedure was implemented a second time; leaf disks were cut from the necrotic local lesions (NLL) and inoculated into *N. longiflora* plants. For the systemic symptoms expressed in the leaves, sections were cut and inoculated in the same way into *N. longiflora* plants. This inoculation was carried out separately on plants, with five replicates per treatment. At 20 dpi, separate inocula were prepared from plants with local and systemic symptoms and inoculated into California bell pepper plants. Five replicates per virus-infected plant were used. Symptoms were monitored to be preserved as a source of inoculum. The serological technique employed was performed according to the methodology of Vasquez-Gutierrez et al. (2023), using monoclonal and polyclonal antibodies specific for PVY (SRA 20001/0096) and ToBRFV (SRA 66800/0096).

Molecular confirmation of PVY and ToBRFV was performed on bell pepper plants by RT-PCR, following the methodology described by Rodríguez et al. (2019). The oligonucleotides were ToBRFV-F (5’-AACCAGAGTCTTCTTCCCTATACTCGGAA-3’) and ToBRFV-R (5’-CTCACCATCTCT-TAATAATCTCCT-3’), designed to amplify a 475-bp fragment. For PVY, the oligonucleotides A-(5’ CATTTGTGCCCAATTGCC-3’) and S6-(5’ GGTGAAGCTAATCATGTCAAC-3’) were used (CNRF, 2020). The resulting amplification products were loaded onto a 1.5% agarose gel.

## 2. Inhibition of ToBRFV and PVY transmission from seed to seedling treated with nanoparticles

The preparation of seven concentrations (50, 100, 200, 400, 600, 800, and 1000 mg L^−1^) for each of the nanoparticles (SiO_2_, Ag, and CeO_2_) was initiated to evaluate their efficacy in preventing viral transmission from seed to seedling. The NP solutions were placed in an ultrasonic bath sonicator (KQ3200DE, Shumei, Cangzhou, China) for 45 min to disperse the particles. The virus inocula (ToBRFV and PVY) were prepared using tissue from symptomatic bell pepper plants; the previously described methodology was employed, and one gram of tissue was macerated in a cold, sterilized mortar with 10 mL of general extraction buffer prepared with: Na2SO3, polyvinylpyrrolidone (PVP), powdered egg (chicken albumin), and Tween 20 in 1X PBST at a concentration of 0.1 M, pH 8.0. The PBST solution was prepared as a phosphate-buffered saline solution with Tween 20.

### 2.1. First part of the experimentation

The experiment was divided into two sections; the first section was carried out as follows: mixed inocula prepared with 800 µL solutions of ToBRFV and PVY, as well as individual inocula (400 µL), were placed in Eppendorf tubes. Immediately, California bell pepper seeds were placed in the tubes and incubated for 24 hours; there were 21 seeds per treatment, with each tube constituting a treatment. The infectious sap was removed and dried on sterile wipes; subsequently, it was placed in Eppendorf tubes with 800 µL of nanoparticles per treatment at the concentrations used. They were incubated for 6 hours at room temperature (15–17 °C) in the dark.

At the end of the incubation period, they were placed on sterile paper towels to dry for 12 hours. Subsequently, they were placed in individual Eppendorf tubes, with seven seeds per tube serving as a replicate, and general extraction buffer was added at a ratio of 1:10 (w/v). Three replicates per treatment were performed. The seeds were vigorously homogenized using an electromechanical homogenizer fitted with sterile wooden sticks as an economical alternative to maceration bags for performing the ELISA technique. Nine interactions (virus, nanoparticles, and NP concentrations) were evaluated, as specified in Table 1. In total, 63 treatments were analyzed, along with four controls.

**Tabla 1.** Severity scale for bell pepper plant leaves.

| Class | Severity | Disease reaction |
| --- | --- | --- |
| 0 | 0 | Healthy |
| 1 | 1.81-10.47 | Very slight |
| 2 | 10.5-28.86 | Minimum severity |
| 3 | 28-30-34.57 | Moderate severity |
| 4 | 30.60-43.33 | Severe |
| 5 | 43.4-51.30 | Very severe |
| 6 | 51.30-64 | Moderately aggressive |
| 7 | 65-72.05 | Aggressive |
| 8 | 73-80.54 | Very aggressive |

**Tabla 2.**
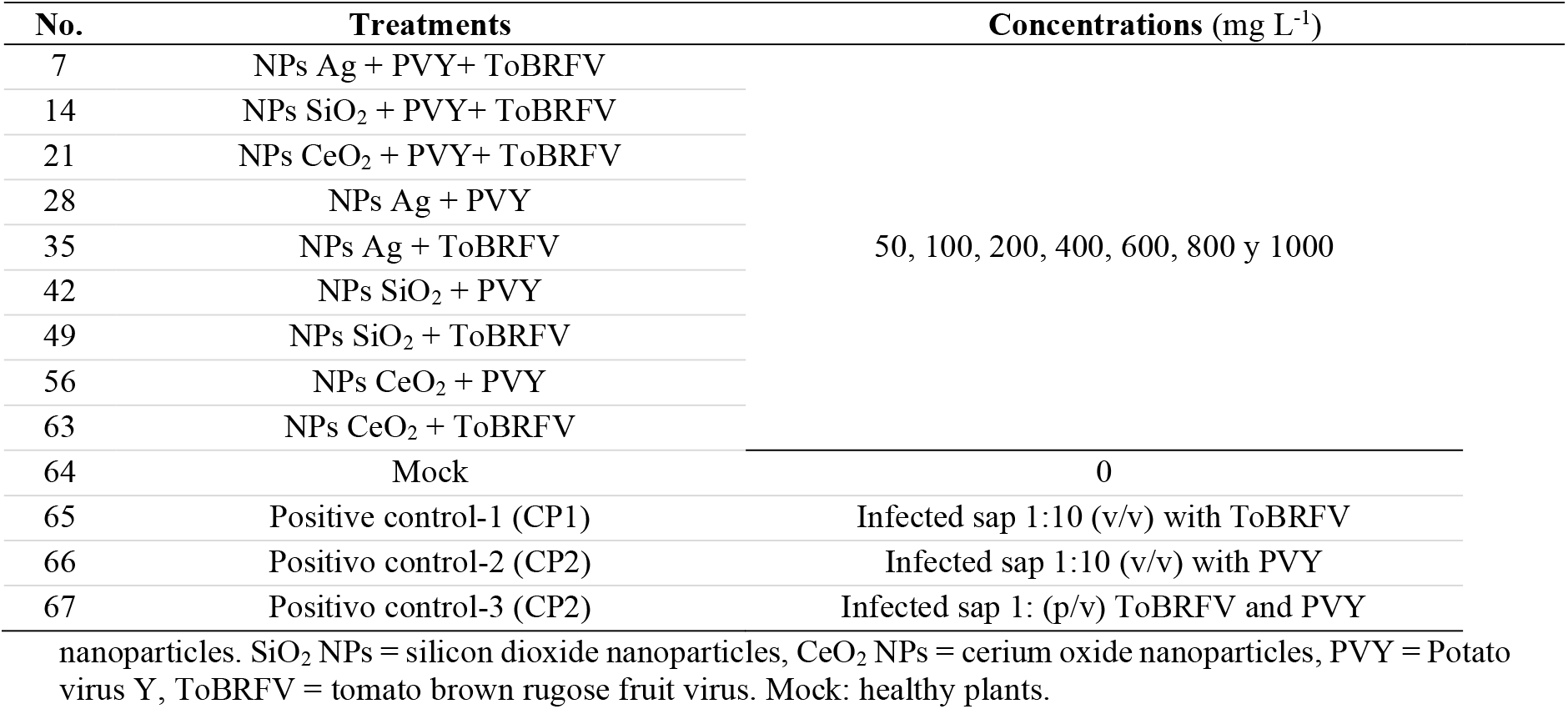
Treatments to evaluate the effectiveness of nanoparticles on seeds and seedlings Ag NPs = Silver.

### 2.2. qELISA assay in seeds

The DAS-ELISA technique (Agdia, Elkhart, USA) began by sensitizing the 96-well plates; 90 µL of capture antibodies per well specific for ToBRFV (ACC 00960) and PVY (Agdia, Elkhart, USA) were added at a 1:200 dilution in 1X Carbonate Coating Buffer (CCB) and incubated (12 h at 6 ± 2 °C) under refrigeration. At the end of the incubation period, the plates were washed three times with 1X PBST buffer. Subsequently, 90 µL of the homogenized seed extract was added to each well, along with the respective controls. The plates were incubated again for 12 h at 6 ± 2 °C, and then washed five times with 1X PBST buffer. Enzyme conjugate for ToBRFV ECA ACC 00960 (Agdia, Elkhart, USA) was prepared at a concentration of 1:200 µL in 1X ECI Buffer (pH 7.5), while for PVY it was a combined solution of detection antibody and enzyme conjugate dissolved in 1X ECM Buffer at a concentration of 1:200 µL (pH 7.5). 90 µL was added to each well, the plate was washed three times, and incubated for 2 h at room temperature. Finally, the PNP substrate was prepared in 1x buffer (pH 9.8) at a concentration of 1 mg/mL, and 90 µL was dispensed into each well of the plate. For all treatments and positive controls, the same technique was used, except for the mock, which consisted of untreated seeds. Incubation was carried out for 30 min, and the optical density (OD) was measured at 405 nm using a Multiskan GO microplate spectrophotometer (Thermo Fisher™, Madrid, Spain), with 5 s of shaking. In total, two readings were taken every 15 min. The viral concentration in the samples was measured based on the average absorbance of the evaluated treatments.

## 3. Second section of the experimentation

The second part began with the addition of the viral inocula to eppendorf tubes. Seed aggregation and treatment of NP concentrations on seeds infected with PVY and ToBRFV were carried out as described in the first section of the experiment for individual and mixed viruses (ToBRFV and PVY). In Eppendorf tubes containing 800 µL of NPs per treatment at the concentrations used, 10 seeds were placed. They were incubated for 6 hours at room temperature (15–17 °C) in the dark. Each treatment was replicated five times, with two seeds per replicate. At the end of the incubation period, they were placed on sterile paper towels for 12 hours. Two seeds were placed in each replicate and sown in 0.5 L polypropylene pots using a 2:1 peat moss and perlite substrate. Five replicates per treatment were used, with each treatment consisting of an NP concentration applied to the virus; the interactions were the same as those employed in the previous experiment (Table 1); there were 63 treatments, three positive controls, and the mock. The seeds were fertigated with a 25% Steiner solution once a week, from emergence until the plants had five true leaves. The environmental conditions under which the seedlings were maintained were 22 ± 2 °C with a photoperiod of 6 to 8 hours of white light per day. Fifteen days after sowing (das), the germination percentage was evaluated. When the plants had four to five true leaves (40 das), they were analyzed by ELISA to assess the viral load in the seedlings. The third section of the leaf was carefully excised, six leaf sections were taken per replicate, and placed in Eppendorf tubes. Two plants per replicate were used for analysis. Six plants per treatment were processed for the quantitative analysis of PVY and ToBRFV, following the ELISA technique described above; vigorous homogenization was performed using an electric homogenizer with sterile pestles. The measurement was performed by optical density (OD) at 405 nm using a Multiskan GO microplate spectrophotometer (Thermo Fisher™, Madrid, Spain). The viral concentration in the samples was measured based on the average absorbance of the evaluated treatments.

### 3.1. Variables evaluated

Incidence and severity were evaluated at 40 dps and calculated using the following formula: I = (IA/N) × 100; where: I = Incidence, which is the percentage of affected individuals (IA) showing symptoms of the disease (%) in a population (N). Severity was calculated using the equation proposed by Vasquez-Gutierrez et al. (2023). The severity was calculated using the equation proposed by Vasquez-Gutierrez et al. (2023). The severity of the disease caused by viral and mixed infections was calculated using the equation: 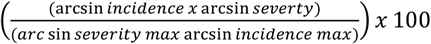, described by Varderplank (1963); Jeger (2004) with modifications, where arc sin represents the arcsine-transformed data, and arc sin severity and arc sin incidence max are the positive control units. To assess the damage caused to seedlings, the quantitative scale for evaluating viral disease in seedlings (Table 1) was used.

Plant height, SPAD index (using a chlorophyll meter, SPAD-502Plus, Minolta), and seed non-germination percentage were evaluated using the equation: PNG = [(Number of ungerminated seeds) / (Number of seeds sown)] × 100 at 40 dps. The viral load in seeds and seedlings, and the disease severity in pepper seedlings inoculated with ToBRFV and PVY, were assessed in a completely randomized design with 63 treatments and five replicates. A mock control and three positive controls were included for each viral species (Table 1). (cuadro 1).

### 3.2. Analysis of phytochemical compounds and radicals

Leaf collection was carried out 50 days after sowing. Apical leaves showing viral symptoms from the evaluated treatments and their respective controls will be selected. The collected leaf samples were stored in a cold cooler until processing. The leaves were washed with 3% NaClO and distilled water. The leaves were placed in Kraft paper bags in a digital drying oven (Felisa FE 291D) at 40°C for 72 hours, until completely dehydrated. Three replicates per treatment were used. The dried samples of both will be homogenized in a Krups GX4100 household grinder and sieved thru a 420 μm mesh; the resulting powder will be stored in amber jars at room temperature (22 ± 2 °C) (Herrera et al., 2022). Extraction will be performed by homogenization using the direct solid–liquid method (Reis et al., 2011). The fine dry powder (20-mesh; ~11.5 g) will be stirred at 25°C with ethanol solvent (C_2_H_6_O) (125 mL) at 150 rpm for 2h and filtered thru Whatman No. 4 paper. The extract solutions were incubated at 2 ± 2 °C for 24 hours. The extracts were filtered to obtain the aqueous phase and placed in test tubes in the dark. The ethanolic extracts were diluted at a 1:10 ratio. Three replicates per treatment were used.

#### 3.2.1. Total phenols

Total phenols were determined using the Folin–Ciocalteu assay. The ethanolic extract solution (1 mL) was mixed with Folin-Ciocalteu reagent (5 mL, previously diluted 1:10 v/v with water) and sodium carbonate (75 g/L, 4 mL). The tubes were shaken for 15 s and allowed to stand for 30 min at 40 °C for color development. Then the absorbance was measured at 725 nm. Gallic acid was used to generate the standard curve (9.4 × 10−3– 0.15 mg/mL), and the results were expressed as mg of gallic acid equivalents (GAE) per L of extract.

#### 3.2.2. Flavonoids

The determination of total flavonoids was carried out using the Dowd method adapted by Arvouet-Grand et al. (1994), which is based on the formation of complexes between flavonoids and 2% aluminum trichloride (AlCl_3_) in ethanol, with absorbance measured at 415 nm. The ethanolic extracts were diluted to keep the values within the spectrophotometric reading range. Quantification was performed by interpolation on a calibration curve constructed with quercetin standard solutions prepared over a concentration range adjusted to the experimental requirements. The results were expressed as milligrams of quercetin equivalents per 100 g of dry weight (mg QE quercetin • 100 g^−1^ DW).

#### 3.3.3. Determination of DPPH and ABTS antioxidant capacity

The antioxidant capacity was determined using the 2,2’-azinobis (3-ethylbenzothiazoline-6-sulfonic acid) cation radical (ABTS+) method, generated by reacting ABTS (7 mM) with potassium persulfate (2.45 mM) and kept in the dark for 12–16 h. The resulting solution was diluted with ethanol to obtain an absorbance of 0.70 ± 0.02 at 734 nm. Subsequently, 2 mL of the ABTS+ solution were mixed with 200 µL of the plant extract, and the reaction was incubated for 6 min in the dark. The decrease in absorbance was recorded at 750 nm using a spectrophotometer, interpolating the obtained values on a calibration curve prepared with Trolox as the standard. The results were expressed as Trolox equivalents per 100 g of dry weight (µmol TE • 100 g^−1^ DW). Antioxidant activity was determined using the 2,2-diphenyl-1-picrylhydrazyl (DPPH) free radical method, prepared in a methanolic solution of known concentration (0.1 mM). Two milliliters of the DPPH solution were mixed with 200 µL of the plant extract and incubated in the dark for 30 min at room temperature. The decrease in absorbance was recorded at 517 nm using a spectrophotometer, comparing the obtained values with a calibration curve prepared with Trolox as the standard. The results were expressed as Trolox equivalents per 100 g of dry weight (µmol TE • 100 g^−1^ DW). The data obtained were subjected to multivariate analysis and Pearson’s correlation (p > 0.05).

## Statistical analysis

To analyze the effect of the treatments on germination and viral concentration, an analysis of variance and mean comparison was performed using Tukey’s multiple range test (P < 0.05). The seed germination percentage data from the evaluated treatments were analyzed using probit analysis to determine the phytotoxic concentration (FC50) for the effect on pepper seed germination. The following equation was applied to the viral concentration results (OD405 nm):

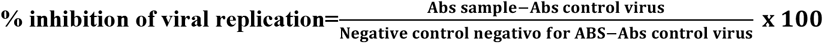

Where: Abs sample: corresponds to the absorbance of the evaluated sample (virus-infected plants treated with NPs); Abs virus control: plants considered as the positive control; Abs negative control: healthy plants considered as the negative control (CN). A Probit analysis was performed, and the inhibitory concentration values (IC_50_ and IC_90_) and their 95% fiducial limits were determined. Using the CF_50_ and CI_50_ data, the selectivity index (SI) was calculated based on the ratio of the 50% phytotoxic concentration to the 50% inhibitory concentration, using the following formula: CF_50_ / CI_50_. The selectivity index (SI) is a parameter used to evaluate an antiviral agent. It is calculated from the ratio between the phytotoxic concentration (FC50) and the concentration that inhibits viral infectivity by 50% (IC_50_); that is, an SI value greater than 20 indicates the antiviral potential of a compound (Chiang et al., 2003). Incidence and severity data, expressed as percentages, were transformed using the arcsine square root for analysis. The variables plant height, chlorophyll (SPAD), disease intensity, phenolic compounds, free radicals, and viral inhibition were subjected to multivariate ANOVA and Pearson’s correlation (P < 0.05). For the analyzes, the statistical software R 4.4.3 will be used. (SAS Institute, 2002).

## Results

### Characterization of nanoparticles

EDAX analysis (Figure 1) reveals the elemental composition profile of the nanoparticle synthesis, suggesting that silver is the primary elemental constituent of the particles. The absorption peak near 3 keV, due to surface plasmon resonance, indicates the presence of Ag NPs (Neira-Vielma et al., 2022). However, signals for C and Si are also observed; these may be due to compounds present in the extract (Vásquez-Gutiérrez et al., 2025v).

**Figure 1.**
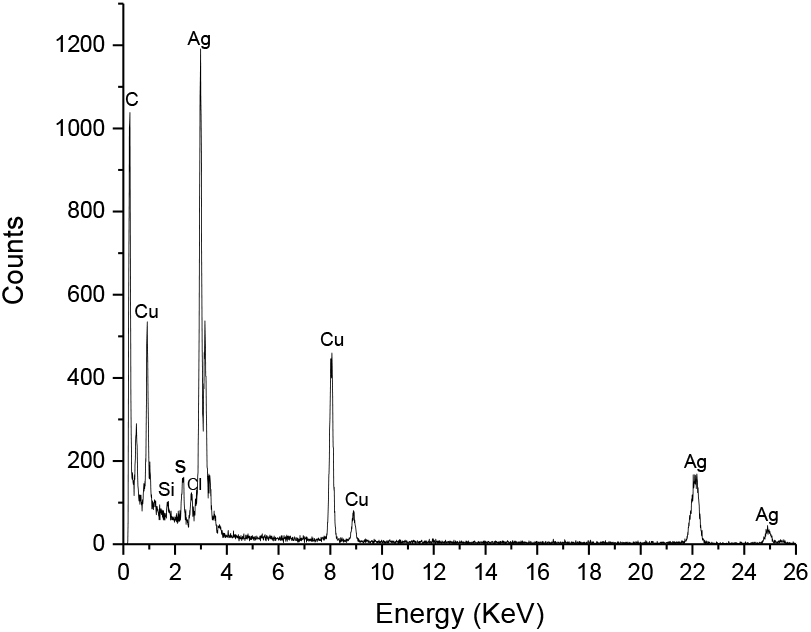
EDX analysis demonstrates that silver (Ag) is the dominant element in the nanoparticles’ composition, while the presence of Cu, Si, and S suggests residual impurities or secondary components from the synthesis process using the extract or the support. For silicon dioxide (SiO_2_) nanoparticles, considering values of K=0.9, *λ* = 0.15406nm, 2*θ* = 22.4°, and instrumental correction, as described above. Suppose: *β*_*meas*_ = 1.0° and *β*_*inst*_ = 0.1° (*Figure* 2*a*).

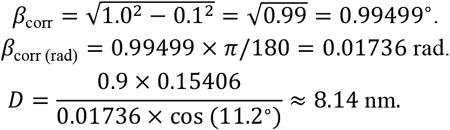

This indicates that the size of the amorphous crystalline domains is 8.14 nm (Figure 2b).

**Figure 2.**
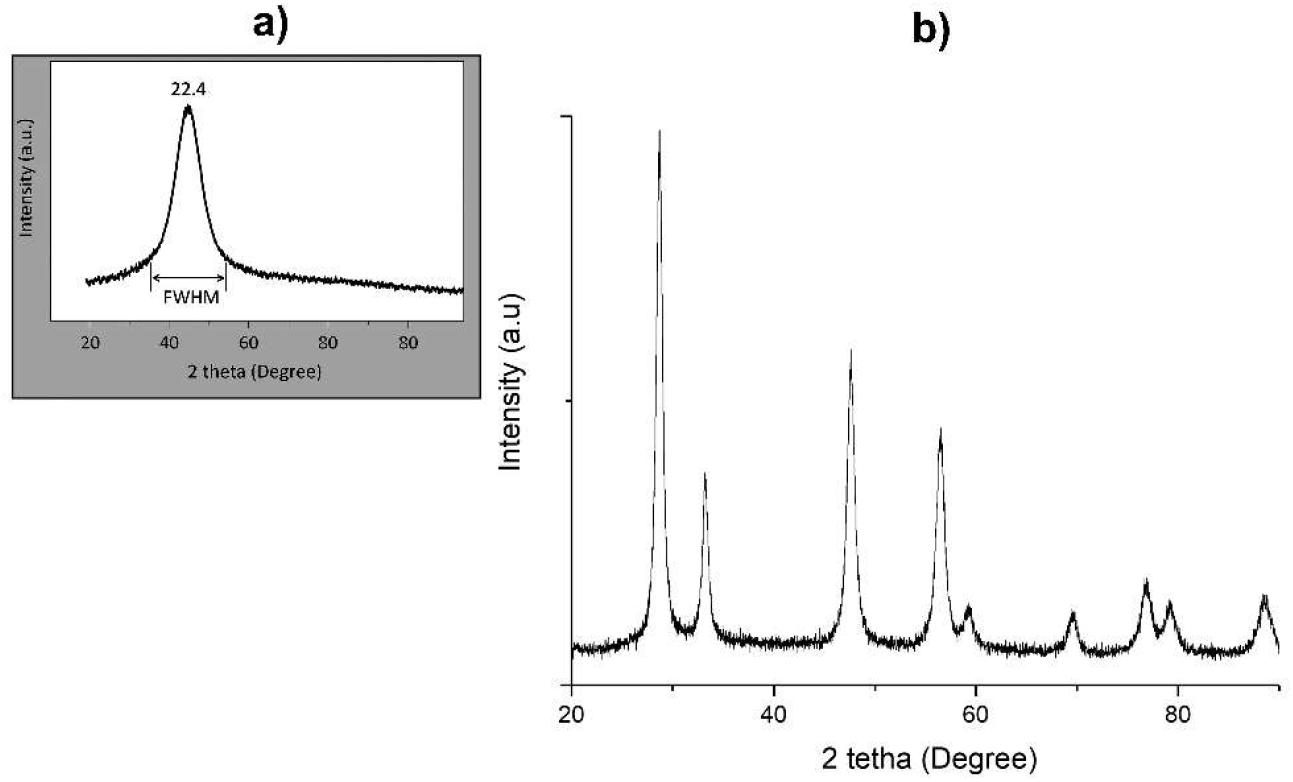
The X-ray diffraction (XRD) pattern of the synthesized SiO_2_ nanoparticles. a) A main diffraction peak is observed at 2θ = 22.4°, corresponding to a semicrystalline structure characteristic of materials with silicon dispersed in or coated by a non-metallic matrix. b) The intensity (a.u.) represents the relative intensity of the diffracted signal, which indicates the number of crystalline planes contributing to the reflection of the X-ray beam. Applying the Scherrer equation to typical halo widths (FWHM ≈ 1.0°) yields estimated crystalline domain sizes of D ≈ 8.14 nm.

The size of the crystalline domains for cerium oxide (CeO_2_) nanoparticles was estimated using the Scherrer equation: 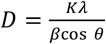. For the analyzed peak (2θ = 21.80°, θ = 10.90°) the measured FWHM was β_*meas*_ = 1.992°. Applying instrumental correction (*β*_*inst*_ = 0.10°) was obtained *β*_*corr*_ = 1.9895°(0.03472 rad). With *K* = 0.9y *λ* = 0.15406nm, the cristal damain size results *D* ≈ 4.07nm (Figura 3).

**Figura 3.**
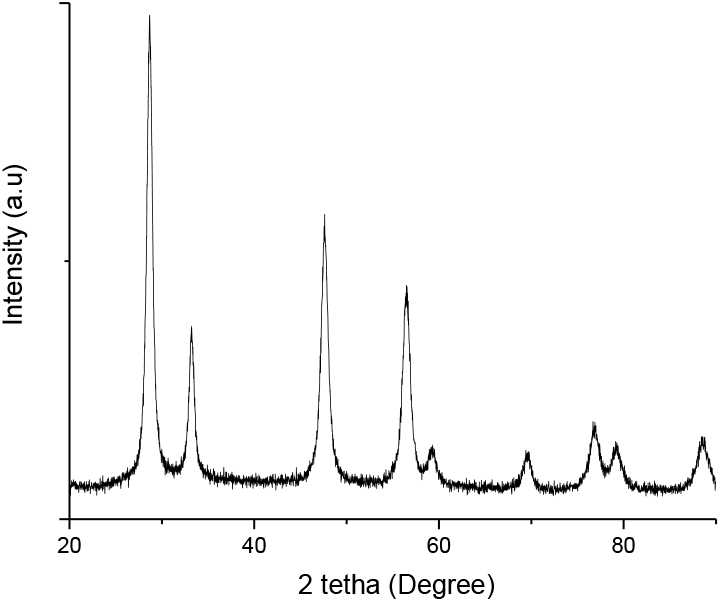
X-ray diffractogram of CeO_2_ nanoparticles obtained by green synthesis. The pattern shows characteristic peaks at 2θ ≈ 28.5°, 33.1°, 47.5°, 56.3°, and 59.1°, corresponding to the (111), (200), (220), (311), and (222) crystal planes of fluorite-type cubic cerium oxide. The main (111) peak was analyzed to estimate the average size of the crystalline domains using the Scherrer equation: 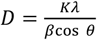. Where *K* = 0.9 (shape constant), *λ* = 0.15406 nm (radiation Cu Kα), *β* it’s the width at half-height (FWHM) corrected for instrumental broadening (*β_corr = √(β_meas*^*2*^ *− β_inst*^*2*^)), and *θ* Bragg angle. From the diffractogram, it was determined β_meas = 1.992° (2θ) at the peak (111), β_inst = 0.1°, it was obtained β_corr = 1.989°. The conversion to radians (β = 0.0347 rad; θ = 14.25°) It allowed estimating an average crystalline domain size of D ≈ 3.9 nm, indicating that the nanoparticles exhibit a finely ordered crystalline structure within the nanometer range. The broadening of the peaks and the absence of significant secondary peaks suggest the high dispersion and phase purity of the synthesized CeO_2_.

For cerium oxide (CeO_2_) nanoparticles, an instrumental correction was applied.

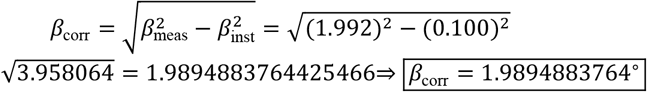

The conversion to radians was performed.

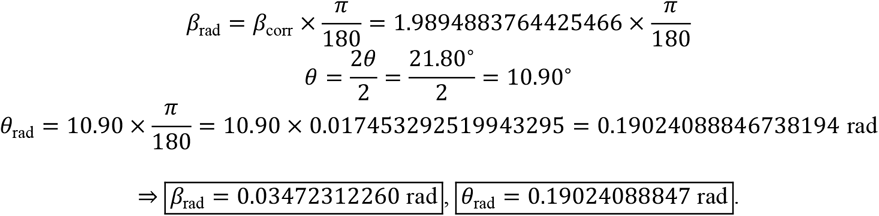

Calculation of the cosine θ

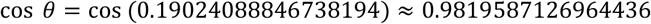

4) Scherrer formula and calculation with corrected β

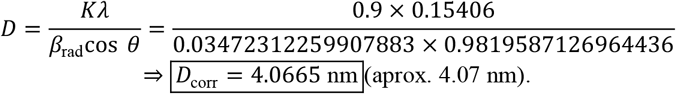

## Molecular characterization of PVY and ToBRFV

Bell pepper plant leaves collected from local fields tested positive for ToBRFV and PVYn using ImnoStrips® (Agdia) (Figure 4 a1 and a2). Tobacco plants inoculated with PVY developed systemic symptoms such as severe mosaic, deformation, and blistering on leaves (Figure 4b). Tobacco plants developed necrotic local lesions (NLL) when inoculated with ToBRFV (Figure 4c). After viral purification, pepper plants inoculated with PVY developed mosaic symptoms and leaf deformation (Figure 4d), while ToBRFV-infected plants exhibited mosaic patterns followed by local necrosis on lower, uninoculated leaves (Figure 4e). The serological test using DAS-ELISA with monoclonal and polyclonal antibodies specific for ToBRFV and PVYn confirmed the presence of both viruses, with absorbances of 1.23 ± 0.3 for PVYn and 1.43 ± 0.4 for ToBRFV, both higher than the negative control (0.011). The PCR products amplified 443-bp fragments for PVY in infected plants (Figure 4f), while for ToBRFV they amplified 475-bp fragments. The results confirm the presence of ToBRFV and PVY in infected plants.

**Figure 4.**
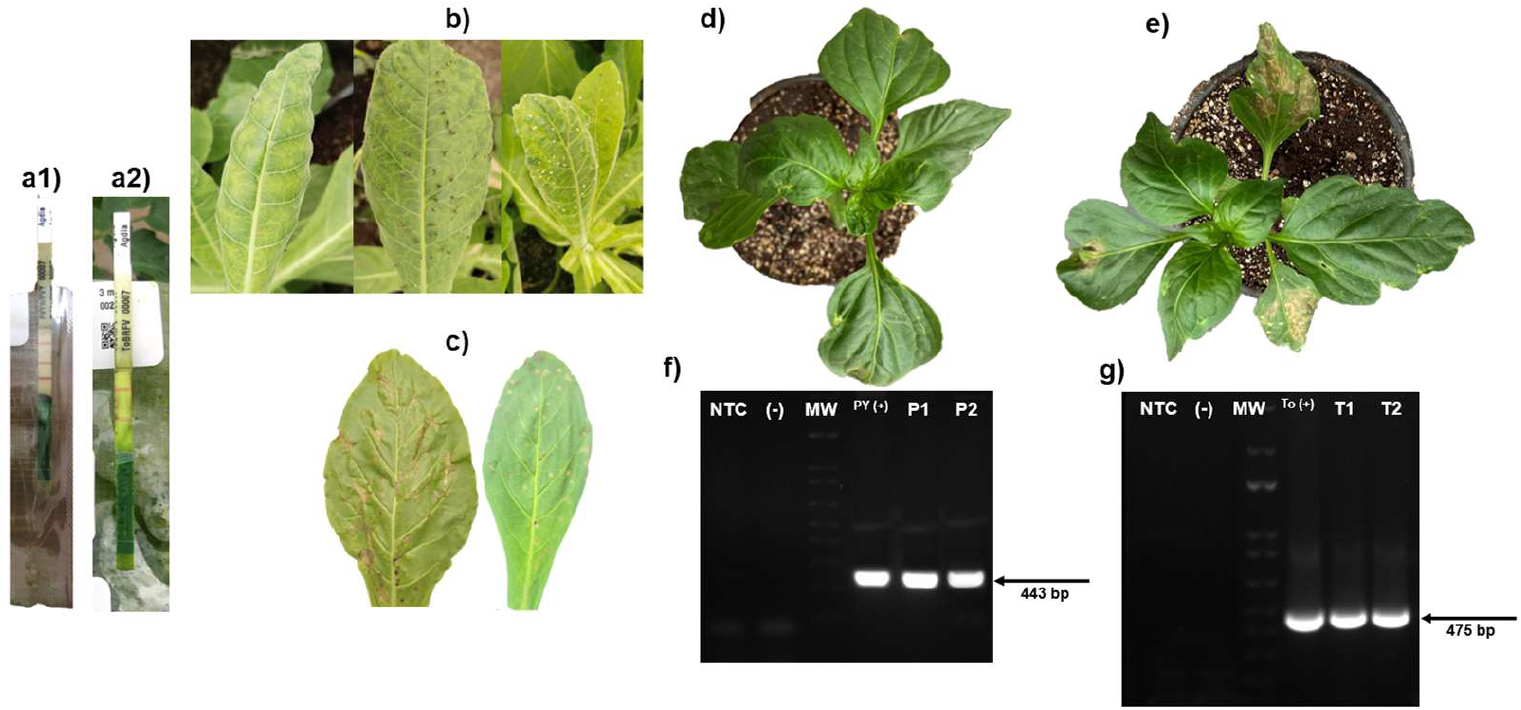
Viral isolation and purification of ToBRFV from bell pepper plants. a1-2) Serological detection of PVY and ToBRFV. b) Systemic symptoms of PVYn expressed in *N. longiflora* plants. c) Local necrotic symptoms on tobacco leaves induced by ToBRFV. d) Bell pepper plant with systemic symptoms caused by PVYn. e) plants with systemic and local symptoms caused by ToBRFV. f) Visualization of the amplification of a 443 bp fragment for PVY. g) amplification of a 475 bp fragment for ToBRFV.

## Impact of green-synthesized nanoparticles on the viral load of PVY and ToBRFV in seeds

### Ag NPs on PVY & ToBRFV mixed infection

The results showed a clear differentiation between the treatments evaluated in the controls (Figure 5a). The positive controls (PC1: ToBRFV, PC2: PVY, and PC3: PVY+ToBRFV) exhibited high levels of infectivity, which validates the sensitivity of the biological system used. The Ag treatments against PVY and ToBRFV (AgPT) were superior to the negative control (NC), showing a tendency to decrease as treatment progressed from T1 (50 mg L^−1^) to T7 (1000 mg L^−1^). Treatments T1 thru T5 (600 mg L^−1^) exhibited partial inhibition, with reductions of 55–70% compared to the positive controls, while T6 (800 mg L^−1^) and T7 achieved reductions of 87–92% relative to the positive control values.

**Figura 5.**
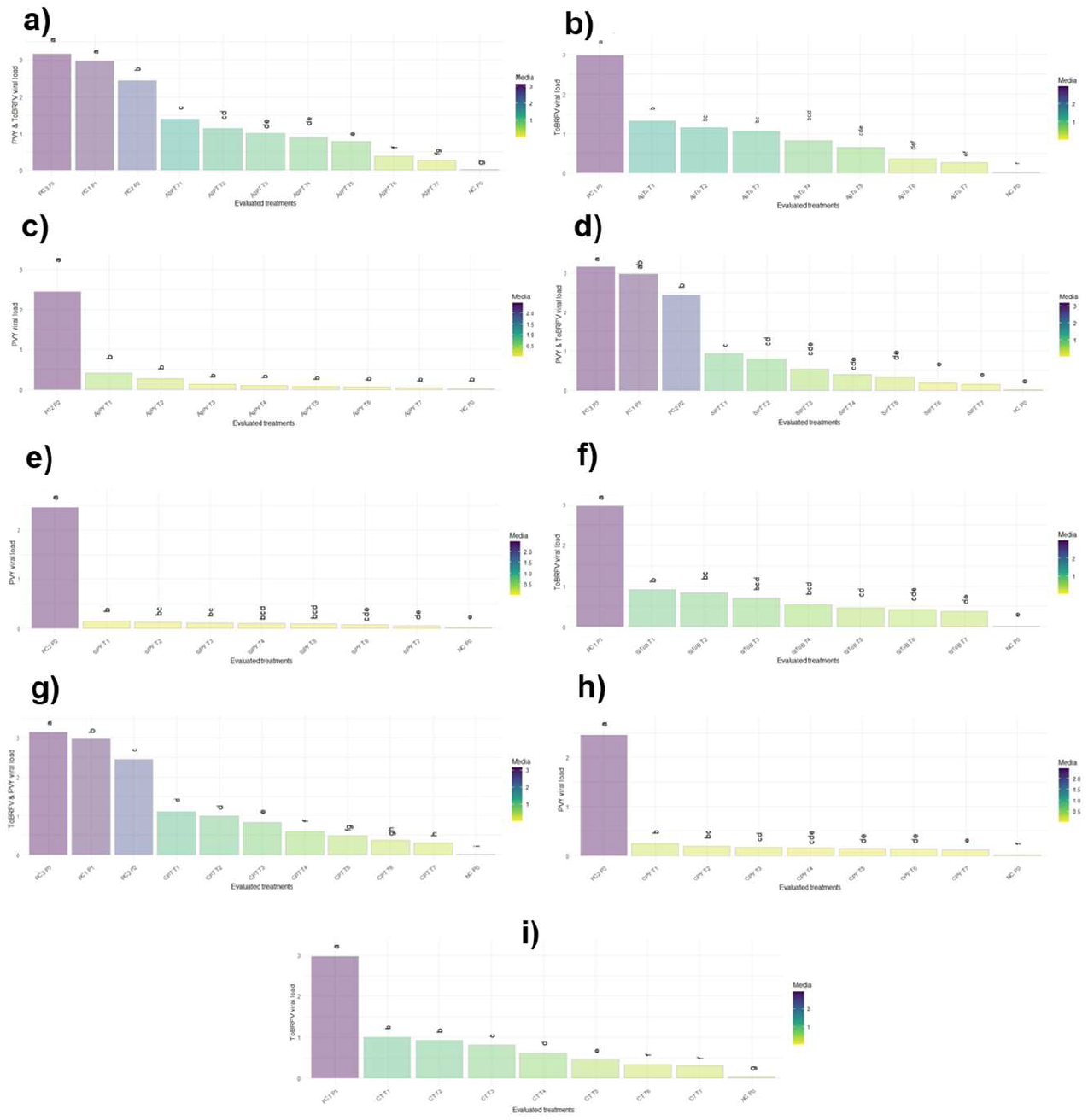
Effect of nanoparticles on the viral titer caused by PVY and ToBRFV co-infection in pepper seeds. PC1 P1: ToBRFV positive control; PC2 P2: PVY control; PC3 P3: mixed PVY and ToBRFV control. The graphs show the interactions between nanoparticles and viral co-infections. a) Silver nanoparticles on PVY and ToBRFV co-infection (AgPT); b) silver nanoparticles on ToBRFV infection (AgTo); c) nanoparticles on PVY infection; d) silicon dioxide nanoparticles on PVY and ToBRFV co-infection (SiPT); e) SiO_2_ nanoparticles on PVY (SiPY); f) SiO_2_ nanoparticles on ToBRFV (SiToB); g) cerium oxide nanoparticles on PVY and ToBRFV co-infection (CPT); h) CeO_2_ nanoparticles on PVY; i) CeO_2_ nanoparticles on ToBRFV (CT). The different nanoparticle concentrations used for each viral interaction were T1: 50; T2: 100; T3: 200; T4: 400; T5: 600; T6: 800; and T7: 1000 mg L^−1^. The ANOVA showed highly significant differences between interactions and treatments (F = 42.34, p < 0.01). Tukey’s multiple comparison test (α = 0.05) grouped the treatments into different letters, indicating significant differences in mean viral load values. Treatment PC3 had the highest mean (3.16 ± 0.19), while NC showed the lowest (0.018). Treatments with different letters differ significantly (p < 0.05).

### AgNPs on ToBRFV

Analysis of the viral load reveals marked differences between treatments T1 (50 mg L^−1^) and T7 (1000 mg L^−1^) with AgNPs on seed infection and the ToBRFV positive control (PC1). PC1 showed an average value of 2.97, which corresponds to maximum infectivity (Figure 5b). In contrast, the treatments exhibited progressive reductions in viral load: T1, T2 (100 mg/L), and T3 (200 mg/L) achieved reductions of 55.9%, 61.2%, and 64.1%, respectively, indicating a moderate antiviral effect. As we progressed to T4 (400 mg/L) and T5 (600 mg/L), inhibition increased to 72.4% and 77.8%, respectively, showing a more pronounced effect. The most effective treatments were T6 and T7 (800–100 mg L^−1^), which achieved reductions of 87.9% and 90.8% compared to PC1, positioning them as the most promising candidates for suppressing ToBRFV viral replication in seeds.

### AgNPs on PVY

The data analysis reveals a significant decrease in viral load in treatments T1–T7 (50 mg/L to 1000 mg/L) compared to the PVY positive control (PC2), whose average value was 2.45 (Figure 5c). Treatment T1 showed an 83.1% reduction, indicating an initial antiviral effect. Subsequently, T2 (100 mg/L) achieved a more pronounced inhibition of 89.2%, while T3 (200 mg/L) reduced the viral load by 95.0%, demonstrating a highly significant effect. Treatments T4 and T5 (400 mg/L and 600 mg/L) further improved this response, with reductions of 96.0% and 96.9%, respectively. Finally, treatments T6 and T7 (800 mg/L and 1000 mg/L) were the most effective, achieving reductions of 97.7% and 98.1% compared to PC2, positioning them as the most promising candidates for practical application.

### SiO_2_ on PVY and ToBRFV

Untreated ToBRFV-infected seeds (PC1) exhibited an average viral load of 2.97 (Figure 5d). Compared to the control, treatment with 50 mg L^−1^ SiO_2_ nanoparticles (T1) resulted in a 68.4% reduction, T2 (100 mg L^−1^) in 72.8%, T3 (200 mg L^−1^) in 82.1%, T4 (400 mg L^−1^) in 86.2%, T5 (600 mg L^−1^) in 88.9%, T6 (800 mg L^−1^) in 93.9%, and T7 (1000 mg L^−1^) in 94.9%. Seeds infected with PVY (PC2) exhibited an average viral load of 2.45. Compared to this value, T1 reduced the viral load by 61.7%, T2 by 66.8%, T3 by 78.2%, T4 by 83.2%, T5 by 86.7%, T6 by 92.6%, and T7 by 93.9%. Finally, seeds infected with ToBRFV and PVY (PC3) reached an average of 3.16. In this scenario, T1 showed a 70.2% reduction, T2 74.4%, T3 83.2%, T4 87.1%, T5 89.6%, T6 94.3%, and T7 95.3%. Overall, the results demonstrate that the treatments were effective against all three viral scenarios. However, relative efficacy varied depending on the virus’s aggressiveness, with more pronounced reduction percentages in the combined control (PC P3) and slightly lower ones against the mild virus (PC P2).

### SiO_2_ NPs on PVY

Seeds infected with PVY (PC2 P2) exhibited an average viral load of 2.45 (Figure 5e). Treatments with silicon dioxide (SiO_2_) nanoparticles on PVY-infected seeds showed very marked reductions in viral load. Treatment T1 achieved a 94.6% reduction, while T2 reduced infectivity by 95.1%. Similarly, T3 achieved a 95.5% reduction and T4 a 96.1% reduction. Treatments T5 and T6 exhibited even greater inhibition, with reductions of 96.6% and 97.0%, respectively. Finally, treatment T7 stood out as the most effective, achieving 98% inhibition compared to PC2 P2.

### SiO_2_ NPs on ToBRFV

Seeds infected with ToBRFV (PC1 P1) exhibited an average viral load of 2.98, while treatments T1–T7 showed much lower values (Figure 5f). Compared to this, all evaluated treatments (T1–T7) showed significant reductions in viral load. Treatment T1 reduced the viral load by 71.8%, while T2 showed a similar decrease of 72.9%. Treatment T3 resulted in a 76.5% reduction, confirming a marked antiviral effect. For their part, T4 achieved 81.5% inhibition and T5 83.5%, highlighting a progressive increase in efficacy. Treatments T6 and T7 were the most effective, with reductions of 84.9% and 87.2%, respectively, compared to PC1 P1.

### CeO_2_ NPs on ToBRFV and PVY

Seeds infected with ToBRFV (PC1 P1) exhibited an average viral load of 2.97 (Figure 5g). In comparison, all treatments showed significant reductions: T1 and T2 (50 and 100 mg L-1) showed reductions of 62.7% and 66%, respectively, while T3 achieved a 71% reduction. For its part, T4 reduced the viral load by 79.1%, followed by T5 at 82.7%. The most effective treatments were T6 and T7, with inhibitions of 87.7% and 89.0%, respectively. When the treatments were compared to the positive control PC2 P2 (PVY), which recorded an average of 2.45, the reductions were slightly smaller but equally noticeable. T1 reduced the viral load by 55.1%, T2 by 58.9%, and T3 by 65.3%. T4 achieved 74.1% inhibition and T5 79.1%, while the most effective treatments were T6 (85.2%) and T7 (87.0%). Finally, compared to the seeds infected with PVY and ToBRFV, they exhibited the highest viral load, with an average of 3.15. T1 reduced the viral load by 64.6%, T2 by 68.3%, and T3 by 73.7%. T4 achieved 81.7% inhibition and T5 84.9%, while treatments T6 and T7 (800 and 1000 mg L-1) were the most outstanding, with reductions of 89.6% and 90.9%, respectively.

### CeO_2_ NPs on PVY

Seeds treated with 50 mg L^−1^ against PVY (CPY T1) recorded a mean of 0.25, corresponding to an 89.8% reduction (Figure 5h). CPY T2 (100 mg L^−1^) reduced the viral load to 0.20, achieving 91.9% inhibition. Similarly, CPY T3 (200 mg L^−1^) showed an average of 0.17, representing a 93.0% reduction. CPY T4 proved even more effective, with an average value of 0.15, equivalent to a 93.8% reduction. For its part, CPY T5 exhibited an average of 0.14, which translates to a 94.5% reduction. CPY T6 maintained consistent results, with an average of 0.13 and 94.8% inhibition. Finally, CPY T7 stood out as the most effective treatment, with an average of 0.11 and a 95.6% reduction in viral load.

### CeO_2_ NPs on TOBRFV

Seeds infected with ToBRFV and treated with 50 mg L^−1^ (CT T1) reached an average value of 1.01, representing a 66.2% reduction (Figure 5i). CT T2 showed a similar average of 0.93, equivalent to a 68.9% reduction. For its part, CT T3 reduced the viral load to 0.81, with 72.7% inhibition. CT T4 showed greater effectiveness, with an average of 0.63 and a 78.9% reduction. Even more outstanding was CT T5, with a mean of 0.46, equivalent to 84.6% inhibition. Treatments T6 and T7 (800 mg/L and 1000 mg/L) stood out as the most effective: the former achieved an average value of 0.33, with an 89.0% reduction, while the latter recorded 0.30, representing 89.9% inhibition compared to the positive control.

### Inhibitory effect of NPs on mixed and individual PVY and ToBRFV infection in seeds

The inhibitory effectiveness of green-synthesized nanoparticles against mixed and individual infections of ToBRFV and PVY showed a considerable reduction (Table 3). Ag nanoparticles against the mixed ToBRFV and PVY infection ranged from 60.4% to 75% (600–1000 mg L^−1^), while SiO_2_ NPs inhibited spread by up to 78.3% and CeO_2_ by up to 72.66%. Ag NPs against ToBRFV infection inhibited viral spread by up to 73.5%, SiO_2_ by up to 69.94%, and CeO_2_ by up to 72.31%. While NPsAg reduced ToBRFV infection in seeds by up to 84%, SiO_2_ reduced it by 48.4%, and CeO_2_ by up to 78.45%.

**Table 3.** Inhibitory effect of green-synthesized nanoparticles on PVY and ToBRFV co-infection in bell pepper seeds.

| Mg L<sup>-1</sup> | Ag NPs<br>PVY&ToBRFV | SiO<sub>2</sub> NPs<br>PVY & ToBRFV | CeO<sub>2</sub> NPs<br>PVY & ToBRFV | Ag NPs<br>PVY | SiO<sub>2</sub> NPs<br>PVY | CeO<sub>2</sub> NPs<br>PVY | Ag NPs<br>ToBRFV | SiO<sub>2</sub> NPs<br>ToBRFV | CeO<sub>2</sub> NPs<br>ToBRFV |
| --- | --- | --- | --- | --- | --- | --- | --- | --- | --- |
| Control | 0±0 e | 0±0 c | 0±0 f | 0±0 b | 0±0 d | 0±0 e | 0±0 c | 0±0 d | 0±0 f |
| 50 | 48.74±3.45 d | 57.34±4.78 b | 53.76±0.44 e | 70.82±19.56 a | 76.79±1.09 c | 72.09±0.98 d | 48.890±2.63 b | 57.08±4.74 c | 54.95±0.56 e |
| 100 | 53.31±0.97 cd | 60.17±6.70 b | 56.21±0.42 de | 72.64±8.22 a | 78.06±1.00 bc | 74.17±1.26 cd | 52.00±2.37 b | 58.62±3.17 bc | 56.46±0.71 e |
| 200 | 55.69±2.19 cd | 66.16±1.67 ab | 59.56±0.95 d | 78.91±4.89 a | 79.08±0.95 bc | 75.47±1.45 bc | 53.71±0.66 b | 61.45±4.28 abc | 59.17±0.58 d |
| 400 | 57.97±2.15 cd | 70.53±9.61 ab | 64.78±2.15 c | 79.70±1.59 a | 79.89±2.74 abc | 76.43±0.35 abc | 58.84±1.74 ab | 65.33±4.76 abc | 63.25±0.63 c |
| 600 | 60.45±3.56 bc | 72.65±8.30 ab | 67.41±0.85 bc | 81.40±1.81 a | 80.23±1.07 abc | 77.39±1.09 ab | 63.69±1.18 ab | 67.63±4.89 abc | 67.55±0.66 b |
| 800 | 70.03±1.24 ab | 77.05±2.06 a | 70.68±2.6 ab | 82.90±0.63 a | 81.9±2.65 ab | 77.8±0.51 ab | 70.89±3.79a | 68.68±2.99 ab | 71.15±0.64 a |
| 1000 | 75.16±9.3 a | 78.29±0.07 a | 72.66±2.33 ab | 84.05±0.41 a | 84.48±2.12 a | 78.45±0.93 a | 73.57±2.80 a | 69.94±2.45 a | 72.31±1.18 a |
| p-value | 0.00414 | 0.00013 | 0.0002 | 0.0606 | 0.0002 | 0.00002 | 0.00359 | 0.0184 | 0.0002 |
|  | CI<sub>50</sub>: 38.44179 | CI<sub>50</sub>: 14.01445 | CI<sub>50</sub>: 18.42881 | CI<sub>50</sub>: 2.69770 | CI<sub>50</sub>: 0.0005636 | CI<sub>50</sub>: 0.0008951 | CI<sub>50</sub>: 36.99811 | CI<sub>50</sub>: 4.47912 | CI<sub>50</sub>: 13.09504 |
| 95% confidence intervals | (3.32811-87.06167) | (4.01201-28.07533) | (4.85886-37.33249) | (0.12286-9.62470) | N/E | N/E | (15.86929-61.13184) | (0.11109-17.46999) | (2.37287-30.29369) |
|  | CI<sub>90</sub>: 1884 | CI<sub>90</sub>: 448.60824 | CI<sub>90</sub>: 1161 | CI<sub>90</sub>: 68.12190 | CI<sub>90</sub>: 2.89566 | CI<sub>90</sub>: 13.80002 | CI<sub>90</sub>: 1547 | CI<sub>90</sub>: 1653 | CI<sub>90</sub>: 1161 |
|  | (797.30584-25810) | (318.23940-736.80763) | (706.07266-2803) | (29.40292-108.59718) | N/E | NE | (944.97614-3543) | (790.49524-10283) | (681.38590-3157) |
| Prediction equation | Y= -1.2016<br>+0.7582(x)<br>CV=7.440977 | Y= -0.9762<br>+0.8514<br>CV=9.010287 | Y= -0.9014<br>Y= 0.7123<br>CV=2.759894 | Y= -0.3939<br>+0.9139 (x)<br>CV=11.26483 | Y=1.1221<br>+0.3454<br>CV=2.433956 | Y= 0.9327<br>+0.3060<br>CV=1.413369 | Y= -1.2395<br>0.7904<br>CV=9.966458 | Y= 0.0871<br>-0.0350<br>CV= 6.683247 | Y= -0.7351<br>0.6580<br>CV=1.241087 |

N/E: Not estimated. CI<sub>50</sub>: 50 % inhibitory concentration; CI<sub>90</sub>: 90 % inhibitory concentration. CV: coefficient of variation. Tukey's multiple comparison test ( $\alpha = 0.05$ ) grouped the treatments with different letters, indicating significant differences in the mean inhibitory effect values. Treatments with different letters differ significantly ( $p < 0.05$ ).

## Impact of green-synthesized nanoparticles on the viral load of PVY and ToBRFV in seedlings

### Ag NPs against ToBRFV and PVY

The ToBRFV positive control (PC1 P1) exhibited an average viral load of 2.98 (Figure 6a). All treatments showed a significant decrease in viral load. Exposure to 50 mg L^−1^ Ag NPs against PVY and ToBRFV infection reduced viral activity by 89.8% (AgPT T1). At 100 mg L^−1^, viral activity was reduced by up to 91.8% (AgPT T2). 200 mg L^−1^ resulted in a 93.3% reduction. Treatments with 400 to 1000 mg L^−1^ inhibited viral spread by 93.9% to 97%. Ag NPs against PVY infection reduced viral spread by 88.3% to 96.7% (50–1000 mg L^−1^). The effect of Ag NPs on mixed infection (PVY and ToBRFV) resulted in 90.6% to 96.7% inhibition of viral concentration (50–1000 mg L^−1^) in bell pepper seedlings.

**Figura 6.**
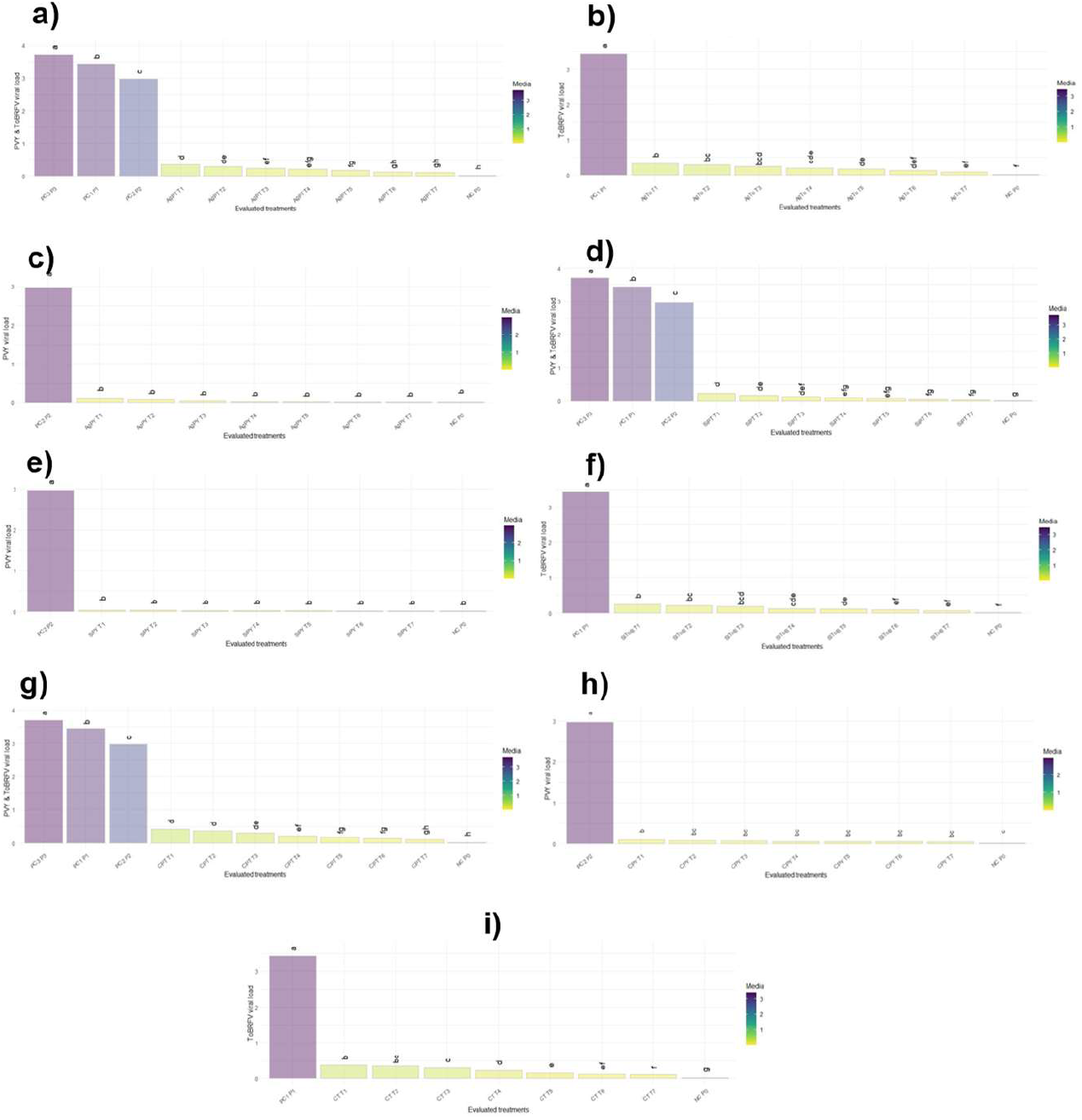
Effect of nanoparticles on the viral titer caused by PVY and ToBRFV coinfection in pepper seedlings. PC1 P1: ToBRFV positive control; PC2 P2: PVY control; PC3 P3: PVY and ToBRFV mixed control. The graphs show the interactions between nanoparticles and viral co-infections. a) Ag nanoparticles on PVY and ToBRFV coinfection (AgPT); b) Ag nanoparticles on ToBRFV infection (AgTo); c) Ag nanoparticles on PVY; d) silicon dioxide nanoparticles on PVY and ToBRFV coinfection (SiPT); e) SiO_2_ nanoparticles on PVY (SiPY); f) SiO_2_ nanoparticles on ToBRFV (SiToB); g) cerium oxide nanoparticles on PVY and ToBRFV coinfection (CPT); h) CeO_2_ nanoparticles on PVY; i) CeO_2_ nanoparticles on ToBRFV (CT). The different nanoparticle concentrations used for each viral interaction were T1: 50; T2: 100; T3: 200; T4: 400; T5: 600; T6: 800; and T7: 1000 mg L^−1^. The ANOVA showed highly significant differences between interactions and treatments (F = 1357.0, p < 0.01). Tukey’s multiple comparison test (α = 0.05) grouped the treatments into different letters, indicating significant differences in mean viral load values. Treatment PC3 had the highest mean (3.70), while NC had the lowest (0.011). Treatments with different letters differ significantly (p < 0.05).

### Ag NPs on ToBRFV

Ag nanoparticles against ToBRFV infection achieved an 89.9% reduction in viral load in pepper seedlings when exposed to 50 mg L^−1^, while AgTo T2 (100 mg L^−1^) achieved a 91.3% reduction (Figure 6b). The 200 mg L^−1^ treatment (AgTo T3) showed an efficacy of 92.2%. The 600 mg L^−1^ (AgTo T5) treatment achieved a 93 % reduction in viral propagation. The AgTo T6 and T7 treatments (800 and 1000 mg L^−1^) were the most outstanding, achieving reductions of 96.0% and 97.3%, respectively, compared to the positive control (PC1 P1).

### Ag NPs on PVY

The treatments (T1–T7) showed drastic reductions in viral titer; however, there were no statistically significant differences among them (Figure 6c). The 50 mg L^−1^ treatment (AgPY T1) reduced the viral load by 97.6% compared to PC2 P2. 100 mg L^−1^ (Ag PVY T2) achieved 97.9% inhibition, while AgPY T3 showed a similar effect with a 98.2% reduction. Treatments T5 and T6 (600 and 800 mg L^−1^) were highly consistent, with reductions of 99.3 and 99.4 %, respectively. Finally, T7 was the most effective, achieving nearly 99.6 % inhibition compared to the positive control.

### SiO_2_ NPs on PVY and ToBRFV

The effect of SiO_2_ NPs persisted even after seed treatment. The viral concentration in ToBRFV-infected seedlings reached a value of 3.44 (Figure 6d). The reduction in viral load in bell pepper seedlings ranged from 93.2 to 99.1% (SiPT T1 and SiPT T7) when concentrations of 50–1000 mg L^−1^ were used. The viral concentration in PVY-infected seedlings ranged from 2.97 (PC2 P2). Treatments using the same concentrations achieved reductions from 92.1% (SiPT T1) to 99.0% (SiPT T7). Seedlings infected with both viruses (PC3-P3) exhibited values of 3.7. Nanoparticle treatments (50–200 mg L^−1^) achieved 92–96% inhibition, while T4 and T5 ranged from 96.6 to 98%, and the most outstanding were T6 (98.5–98.8%) and T7 (98.9–99.2%), which achieved almost total suppression of the viral load compared to the PC3-P3 control.

### SiO_2_NPs on PVY

SiPY T1 treatment (50 mg L^−1^) achieved 98.75% inhibition, while SiPY T2 (100 mg L^−1^) reduced the viral load by 98.96% (Figure 6e). Nanoparticles at a concentration of 600 mg L^−1^ (SiPY T5) achieved a 99.27% reduction, followed by SiPY T6 at 99.39%, and finally SiPY T7 (1000 mg L^−1^), which stood out as the most efficient with 99.52% inhibition compared to the positive control. All treatments achieved reductions of over 98.7%, confirming a highly significant effect on PVY suppression in seedlings.

### SiO_2_ NPs on ToBRFV

The treatments exhibited reductions in viral spread compared to the positive control (PC1 P1), which averaged 3.43 (Figure 6f). The 50 mg L^−1^ (SiToB T1) treatment achieved a 93.7% reduction, while SiToB at 100 mg L^−1^ showed a similar effect with a 93.9% reduction. For their part, seeds exposed to 600 mg L^−1^ exhibited 96.7% inhibition, while SiToB T6 (100 mg L^−1^) increased it to 97.2%. Finally, the 100 mg L^−1^ treatment was the most effective, achieving a 98.1% reduction in viral propagation.

### CeO_2_ NPs on PVY and ToBRFV

CeO_2_ NPs treatments (50 mg L^−1^) reduced the viral load by 88.7% for CPT T1, 90.2% for CPT T2, and 91.6% for CPT T3, reflecting progressive inhibition (Figure 6g). CPT T4 and T5 treatments achieved reductions of 94.1% and 95.1%, respectively. The most outstanding treatments were CPT T6 and T7 (800–1000 mg L^−1^), which achieved reductions of approximately 96.7–97.5%, demonstrating the highest antiviral efficacy. It is noteworthy that against the viral combination (PC3 P3), the most severe scenario, T6 and T7 maintain suppression above 96%, positioning themselves as the most effective treatments under coinfection conditions.

### CeO_2_ NPs on PVY

The 50 mg L^−1^ treatment (CPY T1) achieved 96.6% inhibition compared to PC2 P2. 100 mg L^−1^ (CPY T2) showed a 97.4% reduction, while 200 mg L-1 exhibited inhibition of approximately 97.6% (Figure 6h). The 600 and 800 mg L^−1^ treatments (T5 and T6) achieved even more pronounced reductions, with inhibitions of 98.3% and 98.4%, respectively. 1000 mg L-1 (CPY T7) was the most effective, with an average reduction of 98.6%, establishing itself as the treatment with the greatest viral suppression against PC2.

### CeO_2_ NPs on ToBRFV

The treatments showed a significant reduction in viral load compared to the positive control (PC1 P1). The 50 mg L^−1^ CeO2 NPs treatment resulted in an 89.0% reduction compared to PC1 P1 (Figure 6i). The 200 and 400 mg L^−1^ treatments (CT T3 and T4) achieved reductions of 91 and 93.2%, respectively. Subsequently, T5 achieved a more pronounced inhibition with a 95.2% reduction (CT T4).

Effectiveness was further enhanced when the 1000 mg L^−1^ treatment (CT T7) proved most efficient, achieving a 96.8% reduction compared to the positive control.

### Inhibition of nanoparticles on mixed and individual PVY and ToBRFV infection in seedlings

Las nanopartículas Green-synthesized nanoparticles inhibited viral spread by up to 89% in bell pepper seedlings under mixed or individual infection (Table 4). Ag NPs inhibited viral spread of PVY and ToBRFV by up to 81%, with a 50% inhibitory concentration (IC_50_ = 0.04 mg L−1). SiO_2_ NPs reduced mixed viral spread by 86%, with an IC_50_ of 0.20 mg L^−1^, while NPsCeO_2_ inhibited it by up to 80.8%, with an IC_50_ of 0.26 mg L^−1^. The inhibition by Ag, CeO_2_, and SiO_2_ nanoparticles reduced PVY viral spread by up to 89.2%, followed by 88.7% and 84.2%, with an IC50 > 0.002 mg L^−1^. However, against ToBRFV, inhibition was less pronounced, with percentages of 81.3, 83.1, and 80.1 and IC_50_ <0.18 mg L^−1^.

**Table 4.**
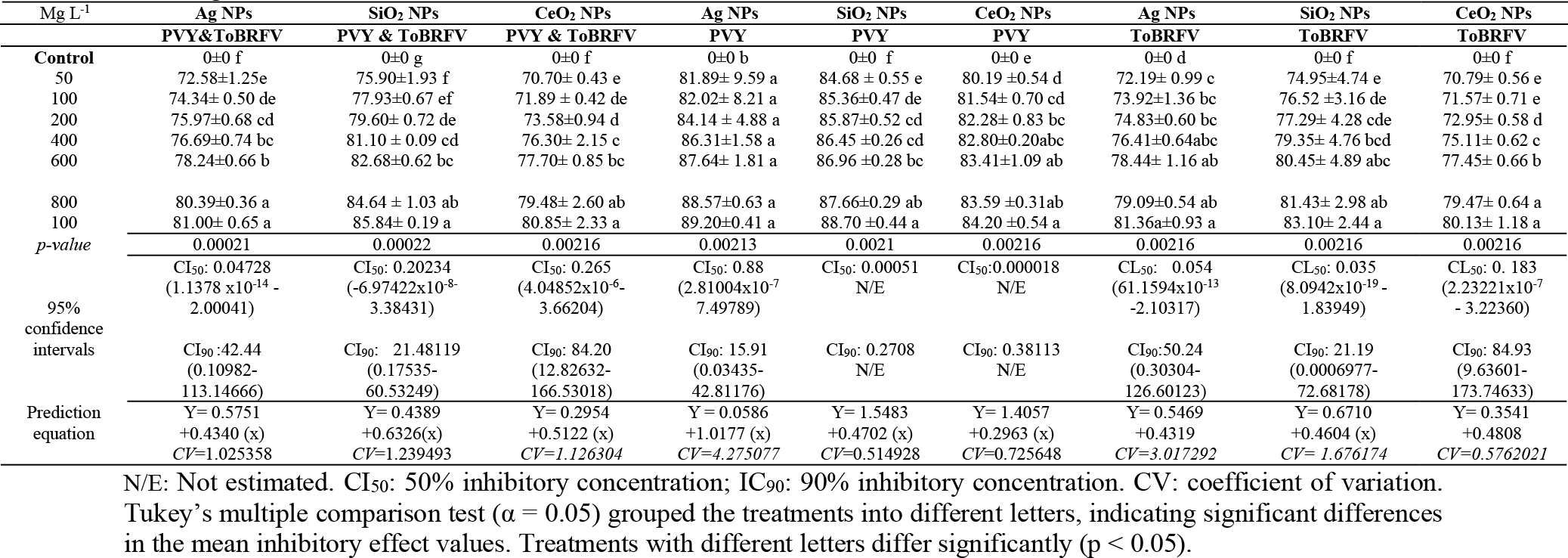
Inhibitory effect of nanoparticles on the viral co-infection of ToBRFV and PVY in bell pepper seedlings.

### Selectivity of nanoparticles with antiviral action

The estimated 50% phytotoxic concentration (EC_50_) of nanoparticles on pepper seed exposure showed high values above 1000 mg L^−1^ to cause phytotoxicity in 50% of the evaluated plant group. The SiO_2_ nanoparticles had an estimated value of 1699 mg L^−1^, compared to Ag NPs, which caused phytotoxicity in plants at 1030 mg L^−1^. The selectivity index (SI) showed values >20, demonstrating antiviral efficacy against both mixed and individual PVY and ToBRFV infections in bell pepper seeds and seedlings. SiO_2_, CeO_2_, and Ag nanoparticles showed a greater antiviral effect against PVY than against ToBRFV. Cerium oxide nanoparticles (CeO_2_) did not cause phytotoxicity in the treatments carried out; therefore, the highest concentration evaluated in this study (1000 mg L^−1^) was taken as the CF_50_ (Table 5).

**Table 5.** Selectivity parameters for nanoparticles with antiviral potential.

|  | Ag NPs<br>PVY & ToBRFV | SiO<sub>2</sub> NPs<br>PVY & ToBRFV | CeO<sub>2</sub> NPs<br>PVY & ToBRFV | Ag NPs<br>PVY | SiO<sub>2</sub> NPs<br>PVY | CeO<sub>2</sub> NPs<br>PVY | Ag NPs<br>ToBRFV | SiO<sub>2</sub> NPs<br>ToBRFV | CeO<sub>2</sub> NPs<br>ToBRFV |
| --- | --- | --- | --- | --- | --- | --- | --- | --- | --- |
| PC<sub>50</sub> | 1030.19 | 1699 | 1000 | 1030.18 | 1699 | 1000 | 1030.188 | 1699 | 1000 |
| 95% confidence intervals | N/E | (1311- 4131) | N/E | N/E | (1311-4131) | N/E | N/E | (1311-4131) | N/E |
| Prediction Equation | -196.318<br>+65.1587 (x) | -15.7691<br>+4.8817 (x) | N/E | -196.318<br>+65.1587(x) | -15.7691<br>+4.8817 | N/E | -196.318<br>+65.1587(x) | -15.7691<br>+4.8817 | N/E |
| P value | 0.0002 | 0.0001 | N/E | 0.01 | 0.0001 | N/E | 0.01 | 0.0001 | N/E |
| IC<sub>50</sub> | 38.44179 | 14.01445 | 18.42881 | 2.69770 | 0.0005636 | 0.0008951 | 36.99811 | 4.47912 | 13.09504 |
| SI | 26.7987 | 121.232 | 54.29 | 381.87 | 3,014,549.3 | 1, 117.19 | 27.844 | 379.32 | 76.37 |

N/E: Not estimated; SI: selectivity index; PC<sub>50</sub>: phytotoxic concentration at 50%; IC<sub>50</sub>: inhibitory concentration at 50%

## Expression of bioactive compounds by viral infection and application of nanoparticles

### Total phenols

The analysis of total phenol contents showed that the treatments exhibited variations compared to the positive controls PC1 P1 (ToBRFV) and PC2 P2 (PVY) (Figure 7). The AgPT T1 treatment stood out with the highest value, exceeding the ToBRFV control by 8% and the PVY control by 19%, suggesting a stimulating effect on phenol accumulation. AgPT T2 treatments remained close to PC1 P1 values (+0.4%) and still exceeded PC2 P2 (+10%), while AgPT T3 reached levels similar to PC2 but 8% lower than PC1 P1. From AgPT T4 onward, phenol accumulation decreased more noticeably, with reductions of 3 to 16% compared to the ToBRFV-infected control, reaching AgPT T7, which showed a reduction of approximately 27%. In plants infected with PVY and treated with 50 mg L^−1^ silver nanoparticles (AgPY T1), the phenol level was only 2–3% lower than the PVY control (PC2 P2). For AgPY T2 (100 mg L^−1^), the values remained unchanged, with a difference of – 1%, suggesting that bell pepper seedlings maintain phenol accumulation at the same level as plants infected with PVY alone. In AgPY T4, the decrease approached –10%. Subsequent treatments showed a more pronounced decrease: PAgPY T5 (600 mg L^−1^) showed a reduction of approximately –12%, followed by AgPY T6 with a decrease of –15%, and finally AgPY T7, which exhibited the lowest accumulation at –20% compared to the PVY control.

**Figure 7.**
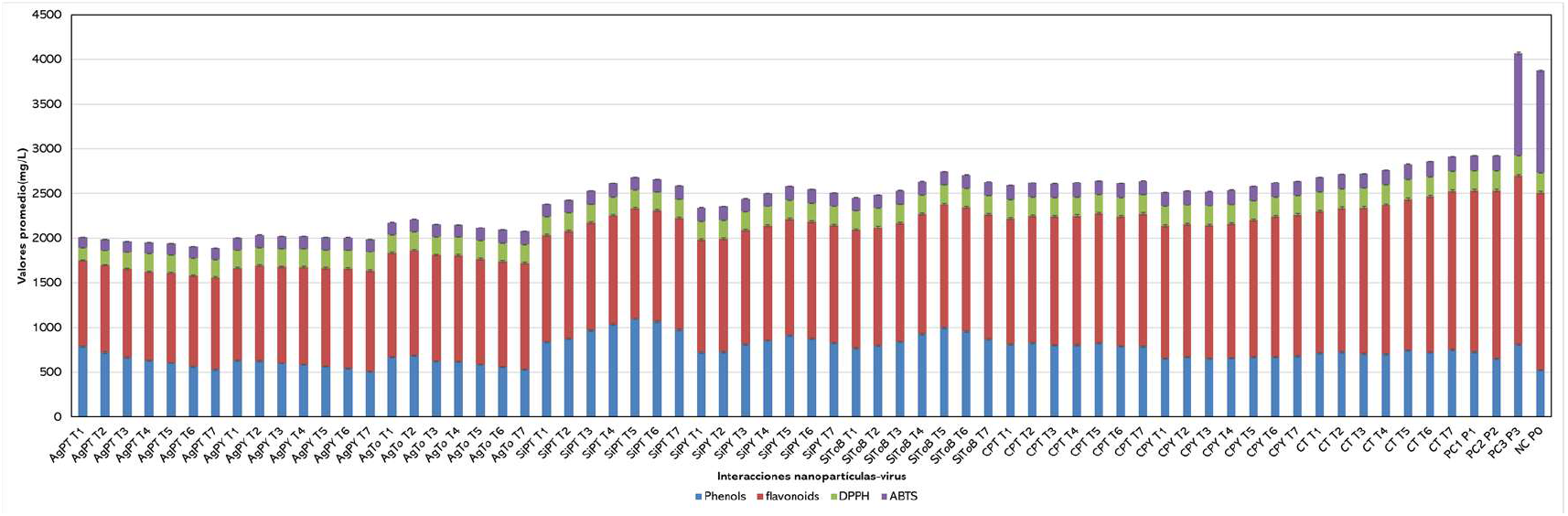
Expression of bioactive compounds in plants treated with nanoparticles and infected with ToBRFV and PVY. Silver nanoparticles on PVY and ToBRFV coinfection (AgPT); silver nanoparticles on ToBRFV infection (AgTo); silver nanoparticles on PVY infection (AgPY); silicon dioxide nanoparticles on PVY and ToBRFV coinfection (SiPT); SiO2 nanoparticles on PVY infection (SiPY); SiO_2_ nanoparticles on ToBRFV infection (SiToB); cerium oxide nanoparticles on PVY and ToBRFV coinfection (CPT). The different concentrations used were T1: 50; T2: 100; T3: 200; T4: 400; T5: 600; T6: 800; and T7: 1000 mg L^−1^. The vertical error bars represent the standard deviations for each treatment.

In AgTo, the initial treatments (T1–T3) maintained values similar to the ToBRFV control, but from T5–T7 significant reductions of up to –25% were observed, indicating a lower capacity for phenolic accumulation in the later stages. In contrast, the T3–T6 (SiPT) treatments achieved increases of +30 to 40% compared to PC2 (ToBRFV). Similarly, in SiPY the intermediate treatments (T2–T5) showed increases of between +18 and 28%, suggesting a response dependent on the application stage. In the case of SiToB, treatments T4–T6 exceeded the control by more than 25%. On the other hand, CPT showed moderate increases, with T3–T5 exhibiting gains of +12 to 20%, while late treatments tended to stabilize without any relevant differences. In contrast, CPY exhibited consistent reductions, particularly in T2–T4, with values of –8 to –12%, indicating lower efficiency. Finally, in CT, treatments T4–T6 showed moderate increases of up to +15%, although lower than those observed in SiPT and SiToB.

### Tota flavonoids

The total flavonoid analysis showed marked differences between treatments and the positive controls (Figure 7). In AgPT, the values were consistent with a decreasing trend as the treatments progressed; while T1 showed levels similar to the PC1 control, from T5–T7 reductions of –18 to –25% were observed compared to the controls, indicating a lower capacity for phenolic accumulation under advanced conditions. SiPT stood out with the highest values: treatments T3–T6 (200 to 800 mg L^−1^) showed significant increases of +35% to +50% compared to the PVY control, peaking at T4–T5, where they even exceeded the ToBRFV control by more than 60%. SiPY showed significant increases in most treatments; particularly T4–T6 (400–800 mg L^−1^), which achieved increases of +25% to +35% compared to PVY. In SiToB, the response was equally remarkable, with treatments T3–T6 exceeding control values by +30 to 45%, and T5–T6 (600–8000 mg L^−1^) being the most outstanding. In CPT, the intermediate treatments (T2–T5) showed moderate increases of +12% to +20% compared to PC2, while the late treatments remained stable. Finally, in plants treated with CeO_2_ NPs (CPY) and infected with PVY, reductions of –10% to –15% were observed in T2–T4 (100–400 mg L^−1^) compared to PVY-infected plants, reflecting a limited effect on flavonoid synthesis. CT exhibited a more stable pattern, with discrete increases in T3–T6 of approximately +8 to 15%, without reaching the magnitude of the increases observed in plants treated with SiO_2_ against PVY and ToBRFV.

### DPPH

Leaf extracts from pepper plants infected with PVY and ToBRFV showed low antioxidant activity, significantly lower than the positive controls (Figure 7). Treatment with 50 mg L^−1^ AgNPs against PVY and ToBRFV resulted in 28.3% inhibition. The values progressively decreased with subsequent treatments (AgPT T2–T7). Extracts from seedlings treated with AgNPs against ToBRFV showed values ranging from 25.2% (50 mg L^−1^) to 14.7% (1000 mg L^−1^). In contrast, extracts from plants treated with SiO_2_ NPs against PVY and ToBRFV (SiPT, SiPY, and SIToB) exhibited higher antioxidant capacity, partially approaching the positive controls. The SiPT T5 treatment (600 mg L^−1^), with 39.98% inhibition. Likewise, SIToB T5 (600 mg L^−1^) exhibited 36.44%. The results show that extracts from treated plants (SiPT, SiPY, SIToB) exhibit greater antioxidant capacity compared to extracts from seedlings treated with CeO_2_ NPs and infected individually or in combination with ToBRFV. However, all evaluated treatments showed significantly lower values than the positive controls PC2 (ToBRFV) and PC3 (PVY and ToBRFV).

### ABTs

In the ABTS assay, the extracts showed a progressive increase in antioxidant capacity reduction (ACR), although they did not exceed the values of the positive controls (PC1, PC2, and PC3) (Figure 7). The highest treatment, AgPT T6 (33.46%), while AgTo T7 (31.07%). In contrast, the sap extracts exhibited higher relative activity: SiPT T7 (41.67%) exceeded the positive control value. Similarly, SiToB T7 (39.96%) compared to CP1. The internal controls showed intermediate values, with CPT T7 (43.36%) standing out by exceeding CP1 by 18.2%. Taken together, these results indicate that, although bell pepper plant extracts exhibit moderate activity, the derivatives from sap infected with PVY and ToBRFV approach or even exceed the ToBRFV and PVY positive controls.

## Impact on the physiological growth and viral damage of bell pepper seedlings

### Plant height

The treatments showed differences compared to the positive controls, demonstrating considerable increases in several experimental groups (Figure 8). The AgTo, SIToB, CPY, and CT treatments exhibited the highest plant height (PH) values, significantly exceeding those of the controls PC1 (6.53), PC2 (7.90), and PC3 (4.33). AgNPs against ToBRFV infection in treatments T6 and T7 (800 and 1000 mg L^−1^) stood out with increases of 15.6% and 23.4%, respectively, compared to the positive control. In SIToB, even more pronounced increases were observed: T5 to T7 (600 to 1000 mg L^−1^) exhibited values of 9.13 to 9.37, equivalent to increases of 15% to 18% compared to PC2, which demonstrates a strong induction of activity. Cerium oxide NPs on PVY showed a progressive upward trend, with T5 to T7 (600 to 1000 mg L^−1^) being the most notable, exhibiting increases from 27.4% (T5) to 52% (T7) compared to PC1, suggesting a stimulated response under combined viral conditions. Similarly, the CT group treatments exhibited the greatest overall differences, with T5 thru T7 (600 to 1000 mg L^−1^) exceeding the positive controls by 34% to 50%, thus establishing themselves as the best-performing treatments for the analyzed variable. In contrast, the AgPT, SiPT, and AgPY groups showed more moderate responses, with increases of less than 10% or similar to the controls, indicating more limited or stable activity against viral factors.

**Figure 8.**
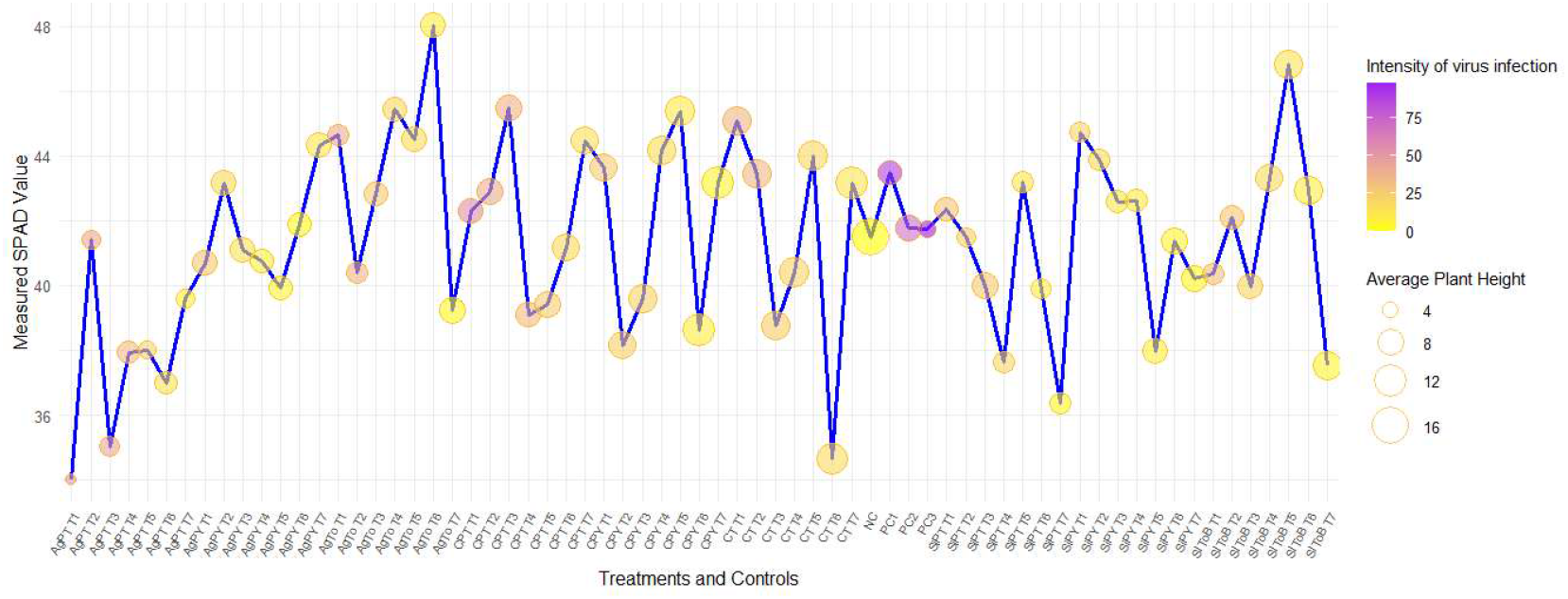
Impact of nanoparticles on the physiological growth and viral damage of bell pepper seedlings. The blue line represents chlorophyll units (SPAD). The intensity of the viral infection is represented by the degree of coloration, where purple indicates a viral load >50, while 0–10 represents a low viral load. The size of the circles represents the average height of the evaluated plants. a) Silver nanoparticles on PVY and ToBRFV coinfection (AgPT); b) silver nanoparticles on ToBRFV infection (AgTo); c) nanoparticles on PVY infection; d) silicon dioxide nanoparticles on PVY and ToBRFV coinfection (SiPT); e) SiO_2_ nanoparticles on PVY (SiPY); f) SiO_2_ nanoparticles on ToBRFV (SiToB); g) cerium oxide nanoparticles on PVY and ToBRFV coinfection (CPT). The different concentrations used were T1: 50; T2: 100; T3: 200; T4: 400; T5: 600; T6: 800; and T7: 1000 mg L^−1^.

### SPAD

SPAD values showed significant variations among treatments compared to the positive controls (PC1 = 43.5, PC2 = 41.77, and PC3 = 41.73) (Figure 8). In general, the AgTo, SIToB, CPY, CPT, and CT treatments showed greater increases, indicating better physiological performance. Within the AgTo group, treatments T4, T5, and T6 stood out with increases of 4.4, 2.3, and 10.4%, respectively, compared to PC2, with AgTo T6 exhibiting the highest value (48.03), indicating a significant improvement in photosynthetic efficiency. In SIToB, treatment T5 showed a 12.1% increase compared to PC2, while in CPY, treatments T4 and T5 also exceeded the reference values with increases of 5.7% and 8.6%, respectively. For their part, CPT T3 (45.47) and CT T1 (45.07) exhibited increases of 8.9 and 7.9% compared to PC2, suggesting a favorable response under the experimental conditions. In contrast, the AgPT and SiPT treatments showed values similar to or slightly lower than the positive controls, with no relevant differences. Overall, the results indicate that the AgTo T6, SIToB T5, and CPY T5 combinations promoted greater stability and accumulation of photosynthetic pigments, representing the most notable responses compared to the positive controls.

### Disease intensity

Treatments with Ag nanoparticles against PVY and ToBRFV reduced the mean intensity as applications progressed (Figure 8). The initial values in the 50 mg L^−1^ AgPY treatments T1 (27.75) and T2 (17.68) were moderate, but from 200 mg L^−1^ AgPY T3 (13.96) onward, a progressive downward trend was observed, with AgPY T6 (2.13) and AgPY T7 (11.27) reaching the lowest values in the group. In relative terms, AgPY T6 reduced the intensity by more than 90% compared to the initial values (T1), reflecting a strong suppression of the infectious process, in contrast to the positive controls. Other treatments showed analogous patterns of decrease, albeit with varying magnitudes. Ag nanoparticles on ToBRFV (AgTo T7, 3.16) and silicon on PVY (SiPY T7, 2.17) exhibited low values comparable to AgPY T6, confirming that the sustained reduction in intensity is consistent across treatments with a high level of biological efficacy. The 1000 mg L^−1^ CeO_2_ nanoparticle treatments against ToBRFV (CPT T7, 14.56) and PVY (CPY T7, 3.78) also showed a notable reduction, with CPY T7 standing out as one of the values closest to the best treatments, exhibiting a decrease of nearly 85%. The AgPY T5–T6 treatments (600 to 800 mg L^−1^), along with SiPY T7 and CPY T7 (1000 mg L^−1^), maintained the lowest intensity levels, suggesting a high capacity to suppress damage or viral replication and superior biological efficiency compared to the other treatments.

## Discussions

Viral concentration data in infected seeds show that some nanoparticle treatments achieved substantial reductions in ToBRFV and PVY viral loads, indicating an antiviral effect at the seedling stage. In particular, intermediate SiPT and SIToB treatments exhibited very low viral loads, reflecting inhibitions of over 90% compared to the positive control, making them promising candidates (Figure 12a, b). At the same time, AgPT treatments at their most aggressive levels (T6 and T7) also showed significant effects, although with greater variability among replicates. These trends suggest that antiviral efficacy depends not only on the nanoparticle (Ag, SiO_2_, CeO_2_) but also on the concentration used (T1–T7). In parallel, in 40-day-old seedlings evaluated for physiological and biochemical variables (phenols, flavonoids, SPAD, height, etc.), it was observed that the most successful treatments in viral suppression also exhibited improvements in defense markers (phenols, flavonoids, plant height, and SPAD). This suggests that nanoparticles not only inhibit viral propagation (Figure 12b), but also induce physiological resistance mechanisms in Capsicum annuum. The use of green-synthesized nanoparticles as agricultural antiviral agents has been documented in some viral species. Silver nanoparticles (AgNPs) have been shown to reduce the accumulation of viruses such as TMV in tobacco (Al-Askar et al., 2023), and cerium oxide nanoparticles (CeO_2_) have been shown to be effective against human viruses such as adenovirus and influenza (Nefedova et al., 2022). It has also been reviewed that ~10 nm AgNPs can interact directly with viral surface proteins, preventing adsorption or entry (Hussain et al., 2022). In plant systems, AgNPs have been used to control viruses in plant crops (Warghane et al., 2024), and antibiotic and antiviral effects have been reported with silica-supported nanoparticles (Sati et al., 2025). Regarding CeO_2_, studies have validated its antiviral activity against human and animal viruses (reduction in viral units (Nefedova et al., 2022), and recent reviews highlight its potential as a metal-based antiviral agent (Zandi et al., 2022). What sets our research apart is the combined application of nanomaterials to PVY and ToBRFV, individually or in co-infection, as well as quantitative measurement (qELISA) in seeds, which few studies have done to date. The successful treatments (SiPT, SIToB, and AgPT) fall within the effective size ranges (approximately 8 nm for SiO_2_ domains and 50 nm for Ag in your system) in which studies have reported antiviral efficacy (AgNPs of approximately 10 nm) (Hussain et al., 2022).

**Figura 12.**
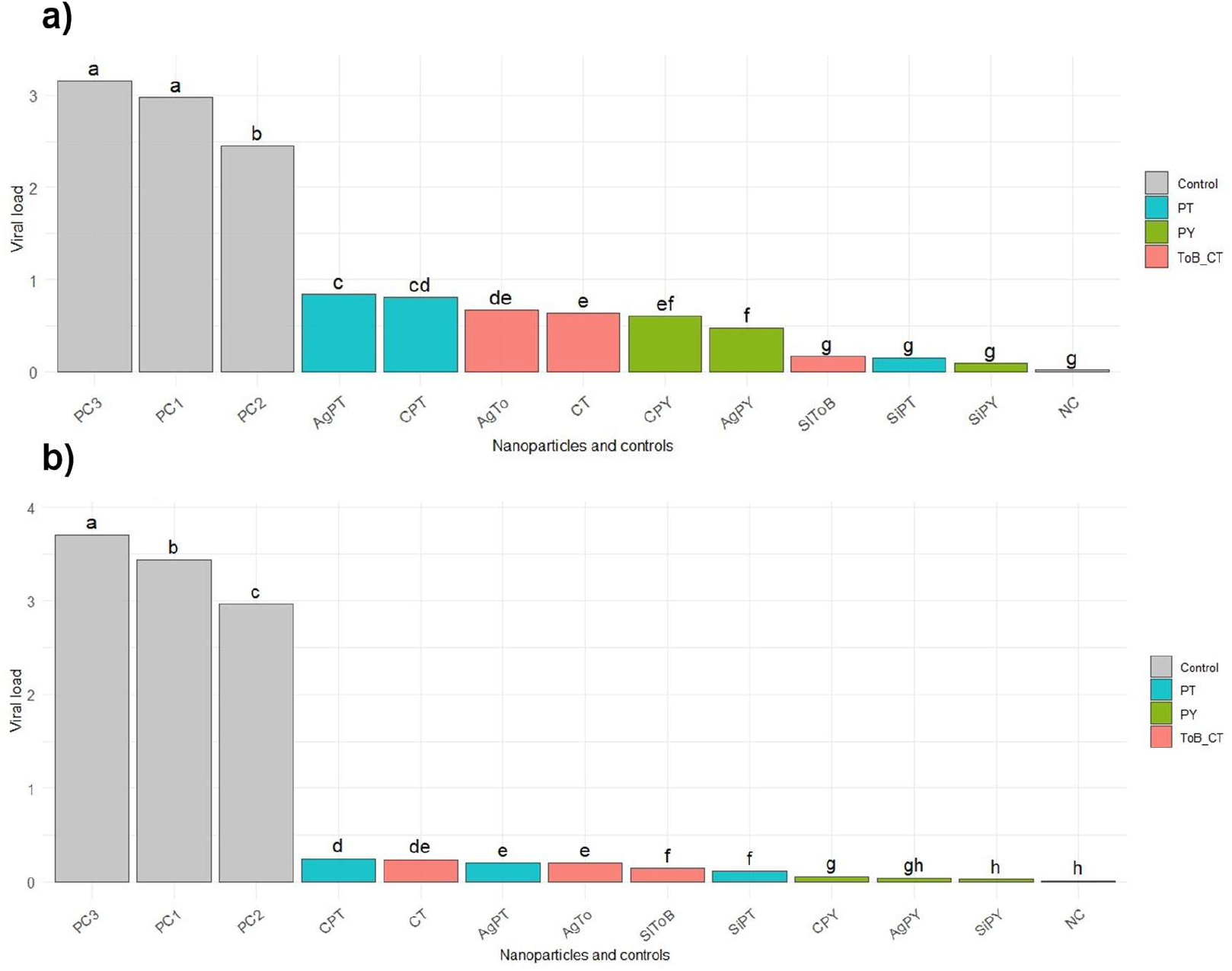
Effectiveness of nanoparticles on mixed and individual ToBRFV infection from seeds to bell pepper seedlings. Effect of nanoparticles (Ag, SiO_2_, CeO_2_) on mixed and individual infection by PVY and ToBRFV, a) in seeds and b) in seedlings. Viral interactions are grouped by color. Treatments with different letters differ significantly (p < 0.05).

The observed efficacy may be due to multiple simultaneous mechanisms. First, nanoparticles can adsorb onto or interact with viral particles or their capsid proteins, preventing the virus from infecting the seed (physical action). Silver nanoparticles, for example, can generate reactive oxygen species (ROS) that damage viral components (proteins, RNA) (Luceri et al., 2023). In your system, the presence of ~50 nm Ag domains provides sufficient contact surface. In the case of SiO_2_ (8.14 nm domain), the nanoparticle can act as a support or vector that enhances the stability of the antiviral agent and promotes controlled release. Furthermore, the induction of plant defense pathways is likely: the increase in phenols and flavonoids in successful treatments suggests activation of the phenylpropanoid pathway, which can reinforce antioxidant defenses and restrict secondary viral replication (Nagai et al., 2015). This is consistent with studies showing that nanoparticles can stimulate the production of antimicrobial secondary metabolites in plants (Vargas-Hernández et al., 2020). The improvement in SPAD, height, and other metrics also indicates that nanoparticles can mitigate viral damage by maintaining photosynthetic integrity and host tissue vigor. The nanometric size, distribution, and surface charge influence penetration and interaction with the plant and virus. In your case, the average values (50 nm for Ag, 8.14 nm for SiO_2_, 4.07 nm for CeO_2_) suggest that the nanoparticles can move efficiently thru seed and seedling tissues, enabling localized action. Among the evaluated treatments, SiPT T3–T6, SIToB T4–T6, and some AgPT T6/T7 emerge as the most successful in suppressing viral load and enhancing biochemical defense. These treatments could be considered as seed treatment protocols to reduce viral load in Capsicum annuum before crop establishment. In particular, the use of silica nanoparticles with curative or protective antiviral activity could offer advantages of lower cytotoxicity compared to pure metals, as well as serving as a microstructural support. The practical application in agriculture could translate into a method for decontaminating infected seeds, complementing conventional phytosanitary control programs with low chemical use. However, it is essential to consider the optimal dosage to avoid phytotoxicity, ensure stability, and minimize synthesis costs. This work is pioneering in applying nanoparticles to ToBRFV- and PVY-infected seeds with quantification by qELISA, but it has limitations: it does not evaluate the possible residual persistence of nanoparticles in harvested plants, nor the exact molecular mechanism of viral inhibition. It would be worthwhile to investigate whether the nanoparticle alone (without added virus) has any residual effect on yield or food safety. As a future perspective, combining fungible nanoparticles with antiviral molecules (RNAi, peptides) could improve efficacy without increasing doses. Furthermore, optimizing the size, charge, and functionalization of nanoparticles (e.g., biodegradable coatings) can enhance selective antiviral action.

The evaluation of the selectivity of green-synthesized nanoparticles against the ToBRFV and PVY viral coinfection revealed significant differences (p < 0.05) in inhibition percentages depending on the type and concentration applied (Tables 3 and 4). Overall, the treatments exhibited a positive dose–response relationship, with inhibition increasing as the nanoparticle concentration was raised from 50 (T1) to 1000 mg L^−1^ (T7). The greatest antiviral effects were observed with silver nanoparticles (AgNPs), achieving inhibitions of 81.0 ± 0.65% for coinfection, 89.2 ± 0.41% against PVY, and 81.36 ± 0.93% against ToBRFV, representing an average increase of ~20–25% compared to SiO_2_ and ~10–12% compared to CeO_2_ at the same concentration (T7). Regarding the median inhibitory concentration (IC_50_), the lowest values were observed for AgNPs, with an IC_50_ of 0.047 mg L^−1^ for coinfection, indicating high antiviral potency and nearly fourfold greater efficacy compared to SiO_2_ (IC_50_ = 0.202 mg L^−1^) and fivefold greater efficacy compared to CeO_2_ (IC_50_ = 0.265 mg L^−1^). These results are consistent with previous observations in which AgNPs directly interfere with the viral capsid or the replicase, inhibiting viral entry or RNA replication (Al-Askar et al., 2023; Mosidze et al., 2025). Additionally, the obtained linear prediction model (Y = 0.5751 + 0.4340x) exhibited a low coefficient of variation (CV = 1.02), confirming experimental consistency. The analysis of the selectivity index (SI), derived from the ratio of the phytotoxic concentration (IF_50_) to the IC_50_, suggests that silver nanoparticles exhibited the highest selectivity (SI > 26.79), followed by SiO_2_ (SI ≈ 121) and CeO_2_ (SI ≈ 76.4). This behavior indicates that Ag nanoparticles are capable of exerting an effective antiviral action at concentrations that do not compromise the host’s physiological development, in contrast to SiO_2_ and CeO_2_, whose therapeutic range was narrower. These results are consistent with the reports by Alowaiesh et al. (2025), who demonstrated greater selectivity of AgNPs against tobacco mosaic virus in solanaceous plants due to their surface interaction with viral proteins and the controlled release of Ag^+^ ions. In single infections, inhibition was consistently higher against PVY (up to ~89%) than against ToBRFV (~81%), suggesting that PVY is more susceptible to nanoparticle–viral protein interactions. This could be attributed to structural differences between the virions or to the variability of movement proteins, as has been described for leek yellow stripe virus-resistant strains (Abdelkader et al., 2025). Finally, the IC_90_ values (15–84 mg L^−1^) confirm that at intermediate concentrations (400–600 mg L^−1^) the nanoparticles reach an antiviral efficacy threshold without affecting germination or plant tissue integrity, validating their potential use as protective or curative agents within an integrated management scheme for viral diseases in bell pepper (Sati et al., 2025; Carrillo et al., 2024).

### Correlation of variables in mixed and individual PVY and ToBRFV infections

Pearson’s correlation analysis established quantitative relationships between biochemical parameters (ABTS radical scavenging activity, DPPH radical scavenging activity, total phenols, flavonoids), physiological parameters (SPAD, plant height), and viral inhibition (Figure 13). Correlation values ranged from r = –0.54 to r = 0.97, with varying levels of significance (p < 0.05, p < 0.01, and p < 0.001) (Hernández et al., 2016). The antioxidant activity measured by RAC_ABTS showed strong positive correlations with total phenolic content (r = 0.81, p < 0.001) and viral inhibition (r = 0.78, p < 0.01), indicating that treatments with higher antioxidant capacity exhibited greater suppression of viral replication. A similar behavior was observed between RAC_DPPH and viral inhibition (r = −0.54, p < 0.01), confirming the consistency of the antioxidant effect on the antiviral response. The disease severity index (DSI) showed significant negative correlations with viral inhibition (r = –0.72, p < 0.001) and with the antioxidant parameter RAC_DPPH (r = 0.60, p < 0.01), demonstrating that the most effective treatments for reducing severity also increased antioxidant capacity. AgPT treatments T1–T3 exhibited weaker or non-significant correlations between phenols and viral inhibition (r = 0.42, p > 0.05), and in some cases negative correlations with RAC_ABTS (r = –0.39, p > 0.05), suggesting a low defense-inducing capacity. In AgPY T5–T7, the correlations between flavonoids and viral inhibition reached r = 0.63 (p < 0.05), indicating a moderate antioxidant response, albeit with intermediate DSI levels. The overall pattern showed that treatments with SiO2 and CeO2 nanoparticles (SiPT, SiPY, CPT) tended to exhibit significant positive correlations between biochemical variables and viral inhibition, whereas silver treatments (AgPT, AgTo) showed weak or negative correlations, associated with a reduced physiological and antiviral response.

**Figure 13.**
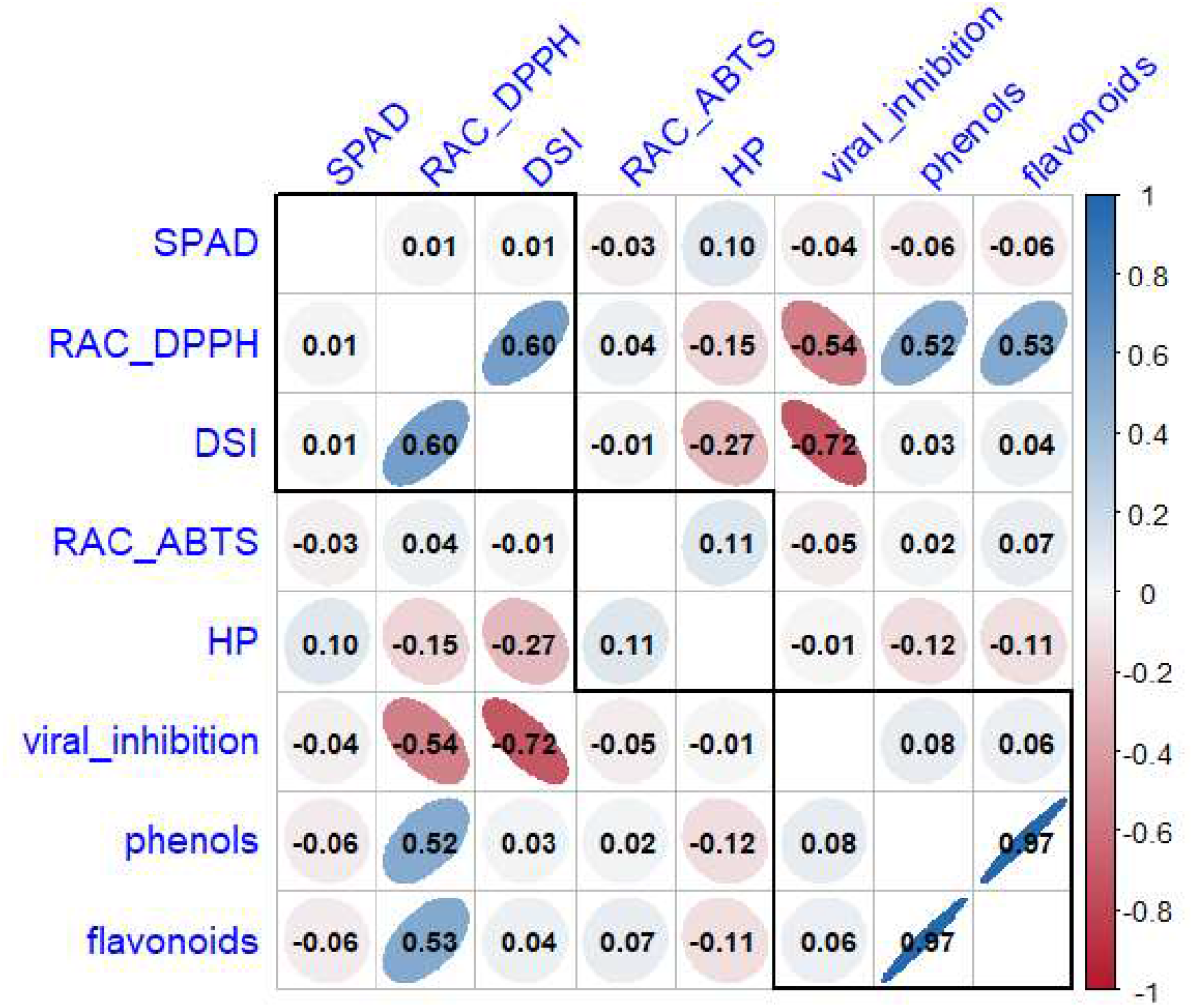
Pearson correlation matrix for agronomic and biochemical variables affected by the mixed viral infection of ToBRFV and PVY, evaluated with nanoparticles. Statistical correlations are shown (p < 0.05). Blue ellipses indicate positive correlations, red ellipses indicate negative correlations, and the degree of tilt reflects the magnitude of the coefficient (r).

### Redundancy analysis of variables affected by the mixed viral infection of ToBRFV and PVY

Redundancy analysis (RDA) showed that canonical axs 1 and 2 jointly explained 56.67% of the total variance observed in the physiological and biochemical data of bell pepper (Capsicum annuum L.) co-infected with PVY and ToBRFV and treated with green-synthesized nanoparticles (Ag, SiO_2_, and CeO_2_). The overall model was significant (p < 0.05 by permutation tests). PC1 explained 33.1% of the variability, while PC2 explained 23.6%, demonstrating a clear structuring of plant responses to the different antiviral treatments. The overall model was significant (p < 0.05) by permutation tests. In the triplot, a clear separation of the treatment groups was observed, with the phytochemical variables related to antioxidant activity—total phenols, flavonoids, and reducing capacity measured by DPPH and ABTS—positively associated with the PC1 axis, showing vectors oriented in the same direction, which indicates a high degree of covariation among these parameters (Figure 14).

**Figure 14.**
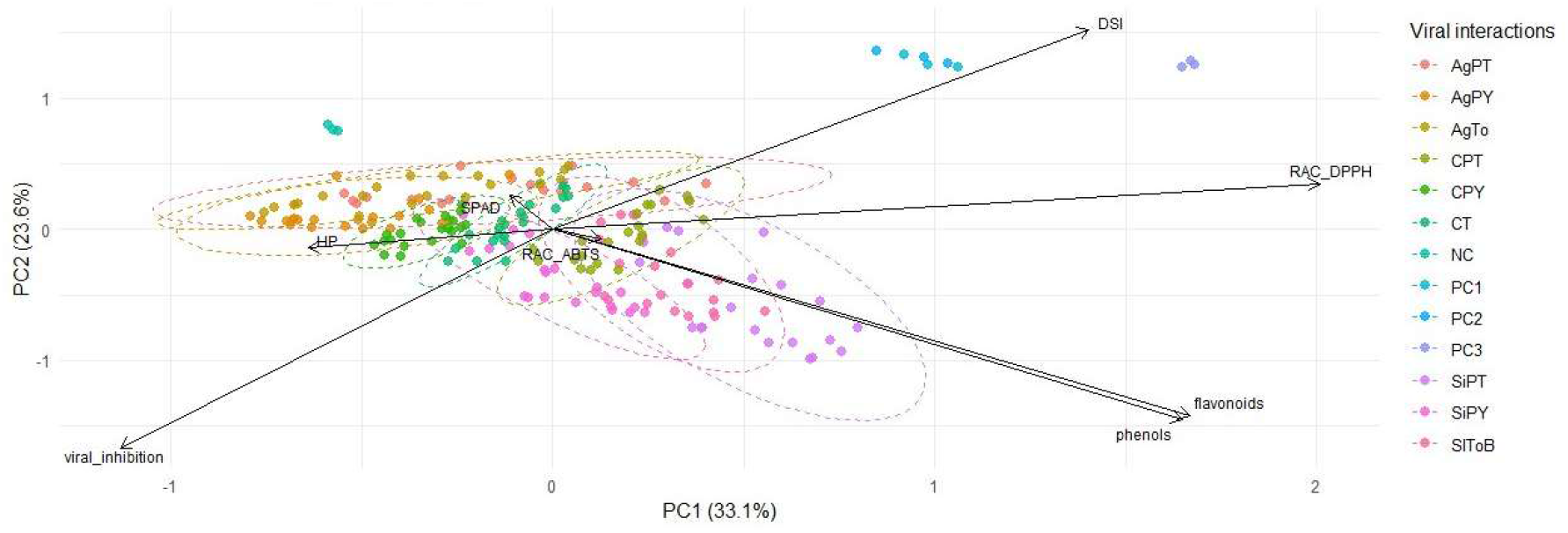
Redundancy Analysis (RDA) of biochemical and physiological variables associated with viral interactions in bell pepper plants. The graph shows the distribution of treatments according to the variation explained by the first two principal component axs (PC1 and PC2), which account for 33.1% and 23.6% of the total variance, respectively. The vectors indicate the direction and magnitude of the variables that contribute most to differentiating the treatments. The ellipses delineate the grouping of treatments according to their general category (AgNPs, SiO_2_NPs, CeO_2_NPs), reflecting the associations between the bioactive compounds and the antiviral response against PVY–ToBRFV mixed infections.

This grouping suggests that nanoparticles, particularly silver (AgNPs) and cerium dioxide (CeO_2_NPs), promoted a coordinated antioxidant response that could contribute to mitigating viral oxidative stress. On the other hand, the physiological variables SPAD (relative chlorophyll content) and HP (plant height) showed negative loadings on PC1, reflecting an opposite behavior to the antioxidant variables, which is consistent with a reduction in oxidative damage and maintenance of the photosynthetic apparatus in the most effective treatments.

The samples corresponding to nanoparticle treatments, especially at higher levels (e.g., T5, T6, T7 at 600–1000 mg/L), are projected in the quadrant closest to the viral inhibition vector, RAC_ABTS, RAC_DPPH, and also relatively close to phenols/flavonoids. This suggests that the treatments achieve a favorable physiological combination: high antioxidant capacity, good phenolic compound content, low damage, and high antiviral efficacy. In contrast, the viral control points (PC1, 2, and 3) cluster toward the DSI vector, indicating that these samples exhibit greater damage, lower antioxidant defense, and reduced (or almost no) viral inhibition. This visually confirms that unprotected viral infection induces a state of stress with greater damage and a weaker defensive response. It is also possible that treatments with different types of nanoparticles (Ag, Si, Ce) form distinct subgroups in the graph: for example, treatments with silica nanoparticles (SiPT) could be positioned more closely aligned with the viral inhibition vector than those with AgPT, suggesting that SiPT has a more potent or consistent effect. In contrast, the DSI (disease severity index) and viral inhibition variables were oriented divergently along PC2: the DSI showed positive correlations, while viral inhibition exhibited negative loadings, indicating that treatments with greater viral inhibition are associated with lower symptom severity. The overall pattern suggests that Ag and CeO_2_ nanoparticles significantly modulated the relationship between antioxidant activity, physiological stress, and viral suppression. This behavior is consistent with previous reports showing that metal nanoparticles increase the expression of antioxidant enzymes and reduce viral accumulation by interfering with virus replication and movement (Alowaiesh et al., 2023; Giménez-Bastida et al., 2021). Although the RDA model did not show any statistically significant constrained components overall, the observed multivariate gradient among the variables indicates that the effects of nanoparticles on antioxidant and viral responses follow biologically relevant trends consistent with the defense mechanisms induced in bell pepper. Taken together, these results reinforce the hypothesis that green nanomaterials can serve as a sustainable alternative for protecting high-value vegetable crops against mixed viral infections (Alharbi et al., 2024).

## Conclusions

The results of this study demonstrate that green-synthesized silver (AgNPs), silicon dioxide (SiO_2_NPs), and cerium dioxide (CeO_2_NPs) nanoparticles exert a notable antiviral effect against single and mixed infections of Potato virus Y and Tomato brown rugose fruit virus in bell pepper. The nanoparticles significantly reduced viral transmission from seed to seedling and viral load in tissues, improving physiological parameters such as the SPAD index and plant height. In particular, CeO_2_NPs exhibited the highest selectivity index (SI > 3000) without any evidence of phytotoxicity, positioning them as safe and highly effective antiviral agents. Pearson’s correlation analysis revealed strong relationships between viral inhibition and antioxidant variables, with total phenolic content and radical scavenging activity (RAC_ABTS and RAC_DPPH) showing significant positive correlations (r > 0.70; p < 0.01). In turn, the disease severity index (DSI) was negatively correlated with viral inhibition (r < –0.80; p < 0.001), confirming that greater antiviral protection is associated with less physiological damage to plant tissue. Additionally, Redundancy Analysis (RDA) explained 56.67% of the total variance, demonstrating that nanoparticles, especially AgNPs and CeO_2_NPs, induce a coordinated antioxidant response that mitigates oxidative stress and promotes antiviral resistance. The variables phenols, flavonoids, ABTS_RAC, and DPPH_RAC clustered with viral inhibition, while DSI showed an opposite pattern, corroborating the efficacy of the treatments at high concentrations (600–1000 mg L^−1^). Taken together, these findings support the notion that the antiviral action of nanoparticles is closely linked to the activation of secondary antioxidant and metabolic defense mechanisms. Green-synthesized nanoparticles, especially CeO_2_NPs, represent a viable and sustainable biotechnological alternative for the preventive management of viral infections in vegetables, contributing to a more resilient agriculture that is less dependent on conventional agrochemicals.

## References

Abdelkader, H.S., Kheder, A.A., Amin, H.A. et al. A comparative study of the antiviral effects of biogenic silver nanoparticles and nanosilica (nSiO2) against Leek yellow stripe virus on Allium sativum L. Eur J Plant Pathol 171, 509–530 (2025). 10.1007/s10658-024-02965-3

Ahmad, M., Ali, A., Ullah, Z., Sher, H., Dai, D. Q., Ali, M., … & Ali, I. (2022). Biosynthesized silver nanoparticles using Polygonatum geminiflorum efficiently control fusarium wilt disease of tomato. Frontiers in Bioengineering and Biotechnology, 10, 988607.

Ahmed, M., Ahmad, S., Abbas, G., Hussain, S., Hoogenboom, G. (2024). Sweet Corn-Bell Pepper System. In: Cropping Systems Modeling Under Changing Climate. Springer, Singapore. 10.1007/978-981-97-0331-9_11

Al-Askar, A. A., Aseel, D. G., El-Gendi, H., Sobhy, S., Samy, M. A., Hamdy, E., El-Messeiry, S., Behiry, S. I., Elbeaino, T., & Abdelkhalek, A. (2023). Antiviral Activity of Biosynthesized Silver Nanoparticles from Pomegranate (Punica granatum L.) Peel Extract against Tobacco Mosaic Virus. Plants (Basel, Switzerland), 12(11), 2103. 10.3390/plants12112103

Alharbi AA, Alharbi M, Alrefaei GI, Aljadani M, Qahl SH, Jaber FA, et al. Valorizing pomegranate wastes by producing functional silver nanoparticles with antioxidant, anticancer, antiviral, and antimicrobial activities and its potential in food preservation. Saudi J Biol Sci. 2024;31(1):103880. 10.1016/j.sjbs.2023.103880

Alowaiesh BF, Alharbi M, Alrefaei GI, Aljadani M, Qahl SH, Jaber FA, et al. Antiviral activity of biosynthesized silver nanoparticles from pomegranate (Punica granatum L.) peel extract against tobacco mosaic virus. Plants. 2023;12(11):2103. 10.3390/plants12112103

Arogundade, O., Ajose, T., Osijo, I., Onyeanusi, H., Matthew, J., & H. Aliyu, T. (2020). Management of Viruses and Viral Diseases of Pepper (Capsicum spp.) in Africa. IntechOpen. doi: 10.5772/intechopen.92266

Arvouet-Grand A., Vennat B., Pourrat A., Legret P., 1994 - Standardization d’une extrait de propolis et identification des principaux constituents, Journal de Pharmacie de Belgique, 49, p. 462–468.

Biswas, T., Guan, Z., & Wu, F. (2018). An Overview of the U.S. Bell Pepper Industry: FE1028, 12/2017. EDIS, 2018(2). 10.32473/edis-fe1028-2017

Carrillo-Lopez, L. M., Villanueva-Verduzco, C., Villanueva-Sánchez, E., Fajardo-Franco, M. L., Aguilar-Tlatelpa, M., Ventura-Aguilar, R. I., & Soto-Hernández, R. M. (2024). Nanomaterials for Plant Disease Diagnosis and Treatment: A Review. Plants, 13(18), 2634. 10.3390/plants13182634

Cognitive Market Research. (n.d.). Bell peppers market report. Retrieved January 8, 2025, from https://www.cognitivemarketresearch.com/bell-peppers-market-report?srsltid=AfmBOopTCtPOkT1JDbIrjR_T0Z81xfiASlcGigpu7FaE6DX7JgEqevUL&utm

Devi, J. et al. (2021). Advances in Breeding Strategies of Bell Pepper (Capsicum annuum L. var. grossum Sendt.). In: Al-Khayri, J.M., Jain, S.M., Johnson, D.V. (eds) Advances in Plant Breeding Strategies: Vegetable Crops. Springer, Cham. 10.1007/978-3-030-66961-4_1

Dwivedi, N., Mishra, M., Sharma, S.S. et al. Genetic analysis and QTLs identification for resistance to the Begomovirus causing pepper leaf curl virus (PepLCV) disease. J. Plant Biochem. Biotechnol. 33, 34–44 (2024). 10.1007/s13562-023-00855-z

Fidan, H., Ulusoy, D., & Albezirgan, H. N. (2024). Exploring Effective Strategies for ToBRFV Management in Tomato Production: Insights into Seed Transmission Dynamics and Innovative Control Approaches. Agriculture, 14(1), 108. 10.3390/agriculture14010108

Giménez-Bastida JA, González-Barrio R, García-Conesa MT, Tomás-Barberán FA, Espín JC. Evidence for health properties of pomegranate juices and extracts. Trends Food Sci Technol. 2021;112:453–467. 10.1016/j.tifs.2021.06.014

Gutiérrez, UV, Treviño, GAF, Ortiz, JCD, Uribe, LAA, Olivas, AF, Beache, MB, & Castillo, FDH (2024). Dióxido de cloro: antiviral que reduce la propagación del ToBRFV en plantas de tomate ( Solanum lycopersicum L.). Virus, 16 (10), 1510. 10.3390/v16101510

Hernández, J. A., Gullner, G., Clemente-Moreno, M. J., Künstler, A., Juhász, C., Díaz-Vivancos, P., & Király, L. (2016). Oxidative stress and antioxidative responses in plant–virus interactions. Physiological and Molecular Plant Pathology, 94, 134–148.

Hussain, F. S., Abro, N. Q., Ahmed, N., Memon, S. Q., & Memon, N. (2022). Nano-antivirals: A comprehensive review. Frontiers in Nanotechnology, 4, 1064615. 10.3389/fnano.2022.1064615

Jeger M. J. (2004). Analysis of disease progress as a basis for evaluating disease management practices. Annual review of phytopathology, 42, 61–82. 10.1146/annurev.phyto.42.040803.140427

Klap, C., Luria, N., Smith, E., Hadad, L., Bakelman, E., Sela, N., Belausov, E., Lachman, O., Leibman, D., & Dombrovsky, A. (2020). Tomato Brown Rugose Fruit Virus Contributes to Enhanced Pepino Mosaic Virus Titers in Tomato Plants. Viruses, 12(8), 879. 10.3390/v12080879

L. C. Chiang, W. Chiang, M. C. Liu, C. C. Lin, In vitro antiviral activities of Caesalpinia pulcherrima and its related flavonoids, Journal of Antimicrobial Chemotherapy, Volume 52, Issue 2, August 2003, Pages 194–198, 10.1093/jac/dkg291

Las sustancias fenólicas endógenas relacionadas con la resistencia de infección por virus. Wang, J., Wang, J., Yue, Z. et al. Disease and Pest Resistance through Phenolic Substances in the Solanaceae. J Plant Growth Regul 43, 2121–2136 (2024). 10.1007/s00344-024-11265-3

Li, N., Yu, C., Yin, Y., Gao, S., Wang, F., Jiao, C., & Yao, M. (2020). Pepper Crop Improvement Against Cucumber Mosaic Virus (CMV): A Review. Frontiers in plant science, 11, 598798. 10.3389/fpls.2020.598798

Liu, S., Tian, W., Liu, Z. et al. Biosynthesis of cupric oxide nanoparticles: its antiviral activities against TMV by directly destroying virion and inducing plant resistance. Phytopathol Res 6, 30 (2024). 10.1186/s42483-024-00250-z

Luceri, A., Francese, R., Lembo, D., Ferraris, M., & Balagna, C. (2023). Silver Nanoparticles: Review of Antiviral Properties, Mechanism of Action and Applications. Microorganisms, 11(3), 629. 10.3390/microorganisms11030629

M.M. Makhsudova, D.T. Jovliyeva, & V.B. Fayziyev (2024). DESCRIPTION OF SOME VIRUSES THAT INFECT THE SWEET PEPPER (CAPSICUM ANNUM) PLANT. Современная биология и генетика, 4 (10), 7–11.

Mosidze, E., Franci, G., Dell’Annunziata, F., Capuano, N., Colella, M., Salzano, F., & Folliero, V. (2025). Silver Nanoparticle-Mediated Antiviral Efficacy against Enveloped Viruses: A Comprehensive Review. Global Challenges, 9(5), 2400380.

Nagai, A., Duarte, L.M.L., Chaves, A.L.R. et al. Potato virus Y infection affects flavonoid profiles of Physalis angulata L. Braz. J. Bot 38, 729–735 (2015). 10.1007/s40415-015-0181-7

Nefedova, A., Rausalu, K., Zusinaite, E. et al. Antiviral efficacy of cerium oxide nanoparticles. Sci Rep 12, 18746 (2022). 10.1038/s41598-022-23465-6

Nefedova, A., Rausalu, K., Zusinaite, E., Vanetsev, A., Rosenberg, M., Koppel, K., Lilla, S., Visnapuu, M., Smits, K., Kisand, V., Tätte, T., & Ivask, A. (2022). Antiviral efficacy of cerium oxide nanoparticles. Scientific reports, 12(1), 18746. 10.1038/s41598-022-23465-6

Neira-Vielma, A.A.; Meléndez-Ortiz, H.I.; García-López, J.I.; Sanchez-Valdes, S.; Cruz-Hernández, M.A.; Rodríguez-González, J.G.; Ramírez-Barrón, S.N. Green Synthesis of Silver Nanoparticles Using Pecan Nut (Carya illinoinensis) Shell Extracts and Evaluation of Their Antimicrobial Activity. Antibiotics 2022, 11, 1150. 10.3390/antibiotics11091150

Ogbole, O.O., Akinleye, T.E., Segun, P.A. et al. In vitro antiviral activity of twenty-seven medicinal plant extracts from Southwest Nigeria against three serotypes of echoviruses. Virol J 15, 110 (2018). 10.1186/s12985-018-1022-7

Ormeño, J., Sepúlveda, P., Rojas, R., & Araya, J. E. (2006). Datura Genus Weeds as an Epidemiological Factor of Alfalfa mosaic virus (AMV), Cucumber mosaic virus (CMV), and Potato virus Y (PVY) on Solanaceus Crops. Agricultura Técnica, 66(4), 333.

Parisi, M., Alioto, D., & Tripodi, P. (2020). Overview of Biotic Stresses in Pepper (Capsicum spp.): Sources of Genetic Resistance, Molecular Breeding and Genomics. International journal of molecular sciences, 21(7), 2587. 10.3390/ijms21072587

Rodríguez-Mendoza, J.; García-Ávila, CJ; López-Buenfil, JA; Araujo-Ruiz, K.; Quezada-Salinas, A.; Cambrón-Crisantos, JM; Ochoa-Martínez, DL Identificación del virus del fruto rugoso marrón del tomate por RT-PCR de una región codificante de la replicasa (RdRP). Rev. Méx. Fitopatol. 2019, 37, 345–356.

Rodríguez-Román, E., León, Y., Perez, Y., Amaya, P., Mejías, A., Montilla, J. O., Ortega, R., Zambrano, K., Olivares, B. O., & Marys, E. (2023). Peppers under Siege: Revealing the Prevalence of Viruses and Discovery of a Novel Potyvirus Species in Venezuela. Sustainability, 15(20), 14825. 10.3390/su152014825

Sánchez-Tovar, M. R., Rivera-Bustamante, R. F., Saavedra-Trejo, D. L., Guevara-González, R. G., & Torres-Pacheco, I. (2025). Mixed Plant Viral Infections: Complementation, Interference and Their Effects, a Review. Agronomy, 15(3), 620. 10.3390/agronomy15030620

Santiago-Meza, Joel de Frías-Treviño, Gustavo Alberto, Aguirre-Uribe, Luis Alberto, & Flores-Olivas, Alberto. (2025). Estimation of losses caused by Potato virus Y in potato crop in Coahuila. Revista mexicana de fitopatología, 43(1).10.18781/r.mex.fit.2404-2

Sati, A., Ranade, T. N., Mali, S. N., Ahmad Yasin, H. K., & Pratap, A. (2025). Silver nanoparticles (AgNPs): comprehensive insights into bio/synthesis, key influencing factors, multifaceted applications, and toxicity─ a 2024 update. ACS omega, 10(8), 7549–7582.

Sati, A., Ranade, T. N., Mali, S. N., Yasin, H. K. A., Samdani, N., Satpute, N. N., Yadav, S., & Pratap, A. P. (2025). Silver Nanoparticles (AgNPs) as Potential Antiviral Agents: Synthesis, Biophysical Properties, Safety, Challenges and Future Directions─Update Review. Molecules, 30(9), 2004.

Secretaría de Agricultura y Desarrollo Rural. (n.d.). México, principal exportador mundial de pimientos frescos: Agricultura. Retrieved January 8, 2025, from https://www.gob.mx/agricultura/prensa/mexico-principal-exportador-mundial-de-pimientos-frescos-agricultura

Senguttuvan, J., Paulsamy, S., & Karthika, K. (2014). Phytochemical analysis and evaluation of leaf and root parts of the medicinal herb, Hypochaeris radicata L. for in vitro antioxidant activities. Asian Pacific journal of tropical biomedicine, 4(Suppl 1), S359–S367. 10.12980/APJTB.4.2014C1030

Shahzadi, S., Fatima, S., Shafiq, Z., & Janjua, M. R. S. A. (2025). A review on green synthesis of silver nanoparticles (SNPs) using plant extracts: a multifaceted approach in photocatalysis, environmental remediation, and biomedicine. RSC advances, 15(5), 3858–3903. 10.1039/D4RA07519F

Shipra Singh, Neelam Sharma. (2023) Indexing of Various Viruses Infecting Capsicum and Their Impact on Its Phytochemical Attributes. International Journal of Biochemistry Research, 8, 1–19

Tatineni, S., & Hein, G. L. (2023). Plant viruses of agricultural importance: Current and future perspectives of virus disease management strategies. Phytopathology®, 113(2), 117–141. 10.1094/PHYTO-05-22-0167-RVW

Tridge. (n.d.). Bell pepper production: Global intelligence. Retrieved January 8, 2025, from https://www.tridge.com/intelligences/bell-pepper/production

Vanderplank, J. E. (1963). Plant diseases: Epidemics and control. New York, NY: Academic Press. https://shop.elsevier.com/books/plant-diseases/van-der-plank/978-0-12-711450-7

Vargas-Hernandez, M., Macias-Bobadilla, I., Guevara-Gonzalez, R. G., Rico-Garcia, E., Ocampo-Velazquez, R. V., Avila-Juarez, L., & Torres-Pacheco, I. (2020). Nanoparticles as Potential Antivirals in Agriculture. Agriculture, 10(10), 444. 10.3390/agriculture10100444

Vásquez GU, Delgado-Ortiz JC, Frías-Treviño GA, Aguirre-Uribe LA and Flores-Olivas A. 2025. Tobamovirusfructirugosum an emergingdisease: review and currentsituation in Mexico. Mexican Journal of Phytopathology 43(1):34.10.18781/R.MEX.FIT.2401-7

Vásquez-Gutiérrez U, López López H, Frías Treviño GA, Delgado Ortiz JC, Flores Olivas A, Aguirre Uribe LA and HernándezJuarez A. 2024. Biological Exploration and Physicochemical Characteristics of Tomato Brown Rugose Fruit Virus in SeveralHost Crops. Agronomy 14 (2): 388. 10.3390/agronomy14020388

Vasquez-Gutierrez, U., Frias-Treviño, G. A., Aguirre-Uribe, L. A., Ramírez-Barrón, S. N., Mendez-Lozano, J., Hernández-Juárez, A., & García-Ruíz, H. (2025v). Nanoparticles and Nanocarriers for Managing Plant Viral Diseases. Plants, 14(20), 3118. 10.3390/plants14203118

Vasquez-Gutierrez, U.; Frias-Treviño, G.A.; Delgado-Ortiz, J.C.; Aguirre-Uribe, L.A.; Flores-Olivas, A. Severity of Tomato Brown Rugose Fruit Virus in tomato (Solanum lycopersicum L.) from a region of Coahuila, México. Int. J. Hortic. Agric. Food Sci. 2023, 7, 1–6.

Visintini Jaime, M. F., Redko, F., Muschietti, L. V., Campos, R. H., Martino, V. S., & Cavallaro, L. V. (2013). In vitro antiviral activity of plant extracts from Asteraceae medicinal plants. Virology journal, 10, 245. 10.1186/1743-422X-10-245

Warghane, A., Saini, R., Shri, M., Andankar, I., Ghosh, D. K., & Chopade, B. A. (2024). Application of nanoparticles for management of plant viral pathogen: current status and future prospects. Virology, 592, 109998. 10.1016/j.virol.2024.109998

Yilmaz, S., & Batuman, O. (2023). Co-Infection of Tomato Brown Rugose Fruit Virus and Pepino Mosaic Virus in Grocery Tomatoes in South Florida: Prevalence and Genomic Diversity. Viruses, 15(12), 2305. 10.3390/v15122305

Zakir, I., Ahmad, S., Haider, S. T.-A., Ahmed, T., Hussain, S., Saleem, M. S., & Khalid, M. F. (2024). Sweet Pepper Farming Strategies in Response to Climate Change: Enhancing Yield and Shelf Life through Planting Time and Cultivar Selection. Sustainability, 16(15), 6338. 10.3390/su16156338

Zandi, M., Hosseini, F., Adli, A. H., Salmanzadeh, S., Behboudi, E., Halvaei, P., … & Abbasi, S. (2022). State-of-the-art cerium nanoparticles as promising agents against human viral infections. Biomedicine & Pharmacotherapy, 156, 113868.

